# Chromatin profiling reveals reorganization of lysine specific demethylase 1 by an oncogenic fusion protein

**DOI:** 10.1101/2020.05.05.079533

**Authors:** Emily R. Theisen, Julia Selich-Anderson, Kyle R. Miller, Jason M. Tanner, Cenny Taslim, Kathleen I. Pishas, Sunil Sharma, Stephen L. Lessnick

## Abstract

Pediatric cancers commonly harbor quiet mutational landscapes and are instead characterized by single driver events such as the mutation of critical chromatin regulators, expression of oncohistones, or expression of oncogenic fusion proteins. These events ultimately promote malignancy through disruption of normal gene regulation and development. The driver protein in Ewing sarcoma, EWS/FLI, is an oncogenic fusion and transcription factor that reshapes the enhancer landscape, resulting in widespread transcriptional dysregulation. Lysine-specific demethylase 1 (LSD1) is a critical functional partner for EWS/FLI as inhibition of LSD1 reverses the transcriptional activity of EWS/FLI. However, how LSD1 participates in fusion-directed epigenomic regulation and aberrant gene activation is unknown. We now show EWS/FLI causes dynamic rearrangement of LSD1 and we uncover a role for LSD1 in gene activation through colocalization at EWS/FLI binding sites throughout the genome. LSD1 is integral to the establishment of Ewing sarcoma super-enhancers at GGAA-microsatellites, which ubiquitously overlap non-microsatellite loci bound by EWS/FLI. Together, we show that EWS/FLI induces widespread changes to LSD1 distribution in a process that impacts the enhancer landscape throughout the genome.

## INTRODUCTION

Ewing sarcoma is an aggressive bone-associated malignancy characterized by the expression of a translocation-derived fusion oncoprotein, most commonly EWS/FLI.^1–4^ The N-terminal portion of the protein is derived from the *EWSR1* gene and comprises a low-complexity intrinsically disordered domain which recruits transcriptional co-regulators.^5–9^ The C-terminal FLI portion of the protein is derived from the *FLI1* gene, which encodes an ETS-family transcription factor.^2^ EWS/FLI contains the ETS DNA-binding domain (DBD) of FLI and functions as an aberrant transcription factor and chromatin regulator, driving global changes in the epigenetic landscape and gene expression, leading to oncogenesis.^5,8–11^

EWS/FLI binds at loci containing the canonical ETS binding motif, 5’-ACCGGAAGTG-3’, with high affinity.^12^ Additionally, the disordered EWS domain confers novel DNA binding properties to the FLI DBD, such that EWS/FLI preferentially binds stretches of repetitive elements with greater than 7 GGAA motifs, called GGAA-microsatellites (GGAA-µsats).^12–14^ Binding of EWS/FLI to GGAA-µsats often results in aberrant activation of nearby genes.^5,8,13^ This occurs through EWS-mediated recruitment of transcriptional and chromatin regulators, like RNA polymerase II^15^ and BAF complexes^5^, and *de novo* assembly of enhancers.^5,8^ Some GGAA-µsats are associated with gene repression, through mechanisms not well understood.^16^ The factors which determine whether an EWS/FLI target is activated or repressed, at both high affinity (HA) sites and GGAA-µsats, are poorly defined.

Disrupting EWS/FLI-mediated gene regulation through direct targeting of EWS/FLI is not yet clinically feasible. As an alternative approach, we previously demonstrated that treatment with the lysine-specific demethylase 1 (LSD1) inhibitor SP2509 reverses the transcriptional activity of EWS/FLI, and the related EWS/ERG fusion, impairing tumor cell growth and viability.^17^ Prior studies suggested that LSD1 was recruited to EWS/FLI-repressed genes as part of the NuRD-LSD1 complex which interacts with the EWS domain ^9^. Having also unexpectedly found that pharmacological blockade of LSD1 disrupts EWS/FLI-mediated gene activation, we predicted that LSD1 recruitment by EWS/FLI is required for chromatin regulation at upregulated targets throughout the genome. Thus, the transcriptional consequences of LSD1 inhibition would recapitulate those of EWS/FLI depletion.

LSD1, also known as *KDM1A*, was the first enzyme discovered that demethylates histone lysine residues.^18^ The flavin adenine dinucleotide (FAD) cofactor-mediated chemistry limits LSD1 to demethylation of mono- and dimethylated lysines.^18^ Histone H3 lysine 4 (H3K4) is the main substrate for LSD1^18^, though LSD1-mediated demethylation of histone H3 lysine 9 (H3K9)^19^ and non-histone proteins, such as p53^20^ and DNMT1^21^, have been reported. LSD1 lacks a DNA-binding domain, and depends upon interacting with other proteins, commonly RCOR1 (CoREST) and histone deacetylases (HDACs), for recruitment to nucleosomes targeted for demethylation.^22^ LSD1 is essential for stem cell function, enhancer decommissioning during differentiation, and transcriptional regulation through modulation of histone lysine methylation levels.^23^ In cancer, LSD1 has been recently described to contribute to aberrant enhancer silencing and transcriptional regulation^24–26^, and overexpression of LSD1 is reported in a wide variety of hematological and solid malignancies^27–31^, including Ewing sarcoma.^32,33^ Elevated expression correlates with aggressive tumor biology and poor prognosis.^30,34,35^

Given that the clinical analog of SP2509, seclidemstat, is now undergoing clinical investigation in Ewing sarcoma (NCT03600649), we wanted to decipher the functional relationship between EWS/FLI and LSD1. In particular, addressing the role for LSD1 in EWS/FLI-mediated gene activation, whether LSD1 contributes to *de novo* enhancer formation, and how EWS/FLI impacts LSD1 genomic localization. In this study we used genomic methods in multiple Ewing sarcoma cell lines, as well as in a specific cell line, A673, either with wildtype expression of EWS/FLI, with EWS/FLI knocked down as a model for a Ewing sarcoma precursor cell, or with EWS/FLI depletion rescued with ectopic expression of the wildtype fusion. By pairing these studies with our previously published transcriptomic data^36^, we evaluated how EWS/FLI impacts LSD1 function in Ewing sarcoma.

## RESULTS

### EWS/FLI and LSD1 colocalize throughout the genome

Previous investigation of LSD1 inhibition in Ewing sarcoma, both pharmacological blockade^17^ and RNAi-mediated depletion^34^, suggested that the LSD1 function is critical for EWS/FLI-mediated gene regulation. We therefore used genomic localization studies to ask whether LSD1 colocalized with EWS/FLI at both activated and repressed targets in Ewing sarcoma cells. In A673 cells, we detected 42673 EWS/FLI peaks and 15202 LSD1 peaks. 12058 LSD1 peaks (79.3%, p=0) were colocalized with EWS/FLI (Figure 1A-C). Overall LSD1 genomic distribution (Figure 1D) was similar to LSD1 distribution when colocalized with EWS/FLI (Supplementary Figure 1A), with a majority of peaks residing in the promoter, intronic, or intergenic regions.

**Figure 1.**
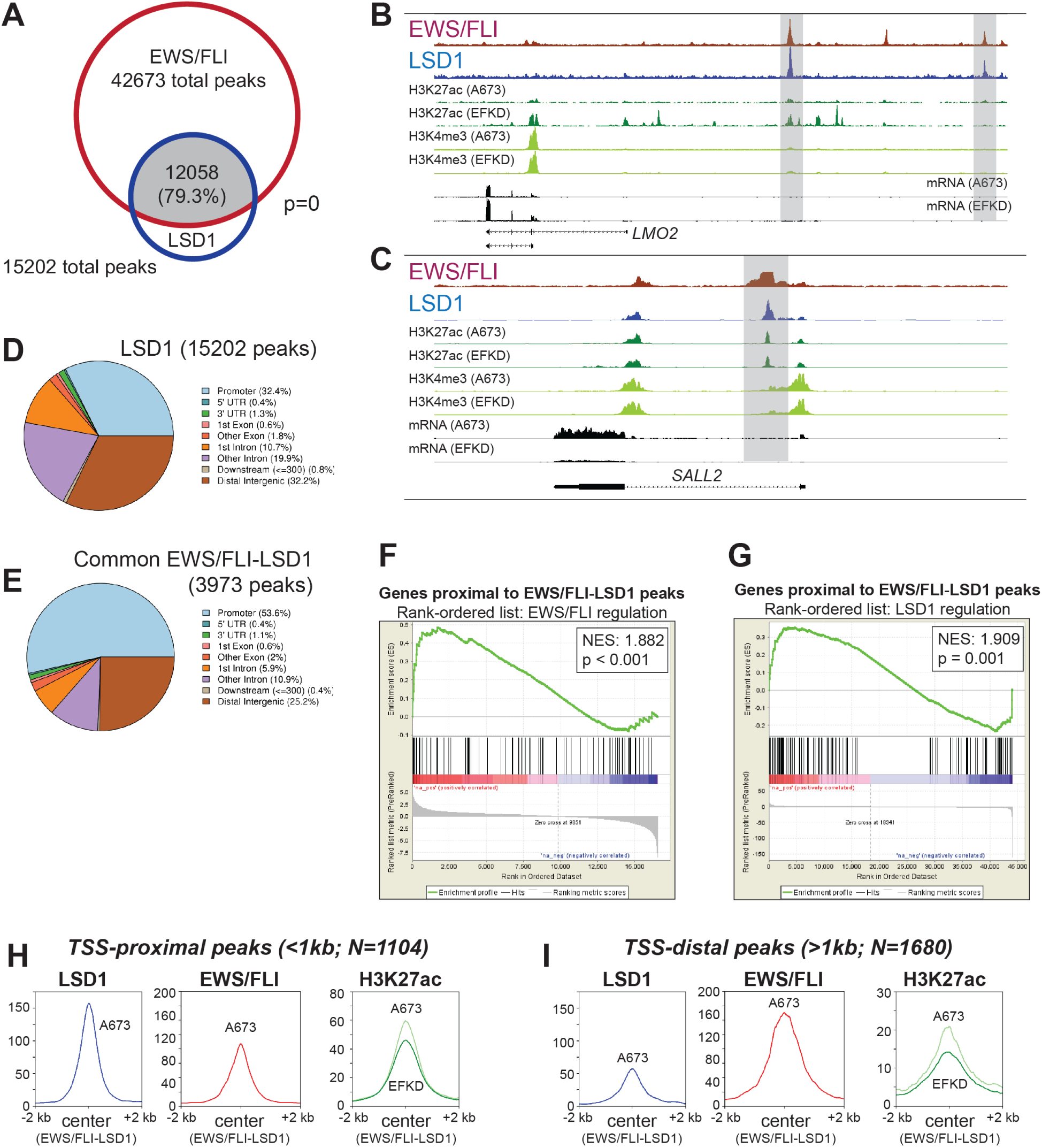
EWS/FLI colocalization with LSD1 is associated with gene activation. A) Venn diagram of EWS/FLI and LSD1 peaks as determined by ChIPPeakAnno; p-value calculated by ChIPPeakAnno. B,C) IGB tracks showing coincidence of EWS/FLI and LSD1 near (B) *LMO2* and (C) *SALL2*. Tracks also show H3K27ac, H3K4me3, and mRNA in the A673 and EFKD conditions. D,E) Genomic distributions of (D) LSD1 peaks in A673 cells and (E) EWS/FLI-LSD1 coincident peaks that are common across all tested cell lines. F,G) GSEA results using promoter-proximal EWS/FLI-LSD1 coincident peaks (<5kb to TSS) as the test set (N = 102) and (F) EWS/FLI gene regulation or (G) LSD1 gene regulation as the rank-ordered dataset. NES=normalized enrichment score. |NES|>1.5 is significant. H,I) Profile plots for signal intensity of LSD1, EWS/FLI, and H3K27ac within 2 kb of EWS/FLI-LSD1 coincident peaks in either A673 cells or EFKD cells as specified. Profile plots are separated into those proximal to (H) or distal to (I) TSS. See also Supplementary Figures 1-7.

These patterns of significant EWS/FLI-LSD1 overlap and LSD1 distribution were consistently observed across additional Ewing sarcoma cell lines: EWS-502, SK-N-MC, and TC-71 (Supplementary Figure 1B-G). We further identified 10300 common EWS/FLI peaks and 6470 common LSD1 peaks that were present in all cell lines (Supplementary Figure 2A-B). Of these 6470 LSD1 peaks, 3973 (61.4%, p=0) showed colocalization with EWS/FLI in all of the tested cell lines, and these displayed a stronger promoter-proximal distribution (Figures 1E, Supplementary Figure 2C-D). Given the large number of EWS/FLI peaks detected here, likely due to the sensitive methodology used for the localization analysis, we re-analyzed the EWS/FLI-LSD1 overlap and distribution in all cell lines using only those peaks with > 8-fold-change in enrichment over background. The resulting overlap and distribution analyses were similar as those described above (Supplementary Figures 3A-E, 4A-E). This set of peaks meeting the higher stringency cutoff was used for all further analyses described below.

Prior studies demonstrated that blockade of LSD1 with the reversible LSD1 inhibitor, SP2509, impaired both EWS/FLI-mediated activation and repression.^17^ In light of this finding we note that LSD1 coincided with EWS/FLI peaks at both activated and repressed target genes in A673 cells (Figure 1B-C, Supplementary Figure 5A-B), suggesting a functional relationship. This was consistently observed in all the assayed cell lines (Supplementary Figure 5C-E). To further explore the function of EWS/FLI-LSD1 colocalization in gene regulation, we next used Gene Set Enrichment Analysis (GSEA)^37^ to evaluate the functional relationship of genes near EWS/FLI-LSD1 co-peaks with either EWS/FLI- or LSD1-mediated gene regulation, previously defined using RNAi-mediated depletion. Genes with EWS/FLI and LSD1 colocalized within 1 kb of transcription start site (TSS) were functionally associated with EWS/FLI-mediated gene activation in all tested cell lines (Supplementary Figure 6A-C). We also found an association between EWS/FLI-LSD1 colocalization and LSD1-mediated gene activation across the tested cell lines (Supplementary Figure 6D-F). The functional association with both EWS/FLI and LSD1 function was likewise observed for common EWS/FLI-LSD1 co-peaks (Supplementary Figure 6G-H). These findings further support a role for LSD1 in EWS/FLI-mediated transcriptional activation in Ewing sarcoma.

We next asked whether EWS/FLI-LSD1 colocalized peaks were also associated with an EWS/FLI-mediated gain of activating histone marks. We examined the levels of H3K27ac, H3K4me1, H3K4me2, and H3K4me3 at loci with EWS/FLI and LSD1 colocalization in A673 cells (wildtype levels of EWS/FLI expression) or cells with EWS/FLI knockdown (EFKD cells). EWS/FLI-LSD1 colocalization was highly associated with increased H3K27ac, consistent with the establishment of a chromatin state which enhances gene activation (Figure 1H-I). Modest increases of uncertain significance were observed for H3K4 mono- and dimethylation, suggesting that LSD1 may not demethylate H3K4 as its primary activity at EWS/FLI-bound loci (Supplementary Figure 7A-B).

### LSD1 is enriched at both GGAA-microsatellites and non-microsatellites

EWS/FLI-mediated gene activation is often modeled as a function of *de novo* enhancer formation following EWS/FLI binding at GGAA-µsats, both proximal and distal to target genes.^5,8,38^ This process involves recruitment of co-activators such as p300^8^ and BAF^5^ by the EWS domain. Given that LSD1-bound regions more strongly associated with gene activation and that inhibition of LSD1 downregulates EWS/FLI-activated genes, we hypothesized that LSD1 would localize to at GGAA-µsats as part of the co-activating machinery assembled by EWS/FLI. We split EWS/FLI-bound loci into GGAA-µsats and non-microsatellites (“non-µsats”) and evaluated EWS/FLI and LSD1 binding at both (Figure 2A, Supplementary Figure 8A-C). While the EWS/FLI binding was stronger at GGAA-µsats than non-µsats, we were surprised that LSD1 was enriched at both GGAA-µsats and non-µsats. The heights of the peaks in the profile plots in Figure 2A and Supplementary Figure 8A-C suggest the amounts of LSD1 colocalized relative to EWS/FLI may be higher at non-µsat loci and LSD1 binding overlapped a majority of EWS/FLI-bound non-µsat regions in all cell lines. The results of HOMER motif analysis for LSD1 further reflected a bias toward localization of LSD1 at non-µsats, as the consensus ETS motif consistently ranked as the most enriched sequence under LSD1 peaks, albeit with GGAA-µsats as the second highest (Figure 2B, Supplementary Figure 8D). These data suggest LSD1 recruitment occurs in a manner distinct from BAF and p300 recruitment as these factors are primarily associated with GGAA-µsat-bound EWS/FLI.

**Figure 2.**
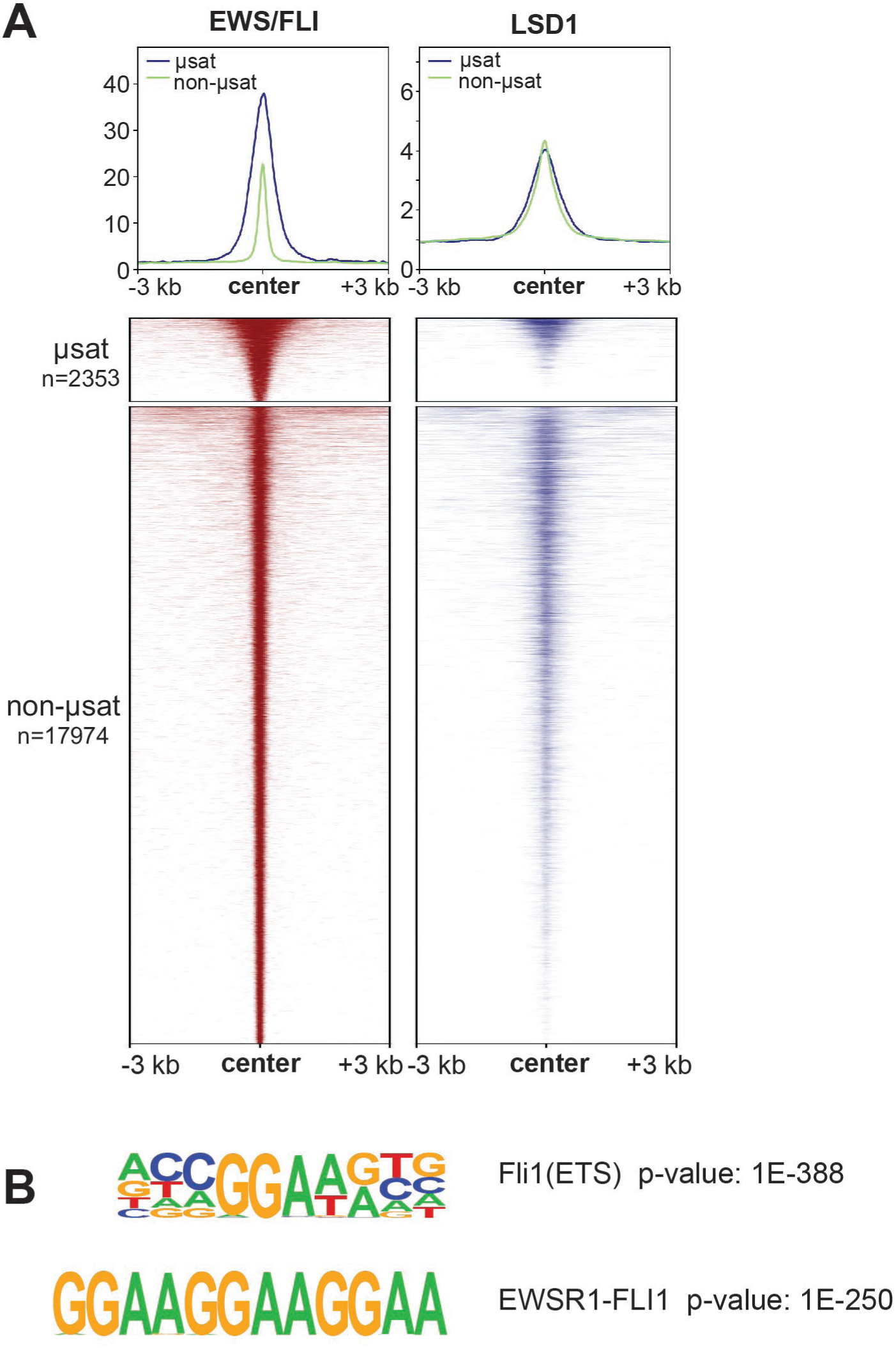
LSD1 is enriched at EWS/FLI binding motifs. A) Profile plots and heatmaps of EWS/FLI (red) and LSD1 (blue) within 3 kb of EWS/FLI peaks. GGAA-microsatellite (µsat) peaks are represented in profile with a blue line and are the top panel in the heatmap. Non-microsatellite (non-µsat) peaks are represented in profile with a green line and are the bottom panel in the heatmap. B) Top ranked result from HOMER *de novo* motif enrichment analysis with significance value. See also Supplementary Figure 8.

### Non-microsatellites play a role in Ewing sarcoma super-enhancers

That LSD1 functions to promote gene activation from both non-µsat and GGAA-µsat loci across the genome is consistent with the previous observation that LSD1 inhibition reverses significant portions of EWS/FLI-mediated transcriptional activity.^17^ We were, however, intrigued by the observations that EWS/FLI-LSD1 sites tended to associate more strongly with non-µsats, as GGAA-µsats are among the most EWS/FLI-responsive elements in Ewing sarcoma. It was unclear how this mechanism fit the model wherein gene activation in Ewing sarcoma is primarily driven by EWS/FLI-mediated changes to the enhancer landscape, through *de novo* deposition of H3K27ac and recruitment of chromatin remodelers to GGAA-µsats.

To further understand the role of LSD1 in EWS/FLI-mediated gene activation, we analyzed the genome-wide relationship between EWS/FLI, LSD1, and the enhancer landscape in A673 cells in more detail. First, we assessed the relationship between super-enhancers (SEs) and EWS/FLI. Using H3K27ac signal overlapping with H3K4me1 signal to define enhancer regions, the Ranked Ordering of Super-Enhancers (ROSE) algorithm^39,40^ identified 833 SEs in A673 cells (Figure 3A, Supplementary Figure 9A; Supplementary Tables 1-2). Previous reports suggest *de novo* enhancers at GGAA-µsats constitute the majority of SEs in A673 cells^41^; we were thus surprised our analysis showed only 20% of SEs overlapped with an EWS/FLI-bound GGAA-µsat (Figure 3A). Instead, the majority (62%) of SEs harbored non-µsat-bound EWS/FLI including 45% that had no GGAA-µsat-bound EWS/FLI and 17% that had both non-µsat- and GGAA-µsat-bound EWS/FLI (Figure 3A). Genes nearest to SEs showed higher levels of expression compared to those near typical enhancers (TEs) (p<0.001; Figure 3B), as expected. Taken together, these data suggest that establishment of super-enhancers at GGAA-µsats may be enhanced through additional EWS/FLI binding at non-µsat sites.

**Figure 3.**
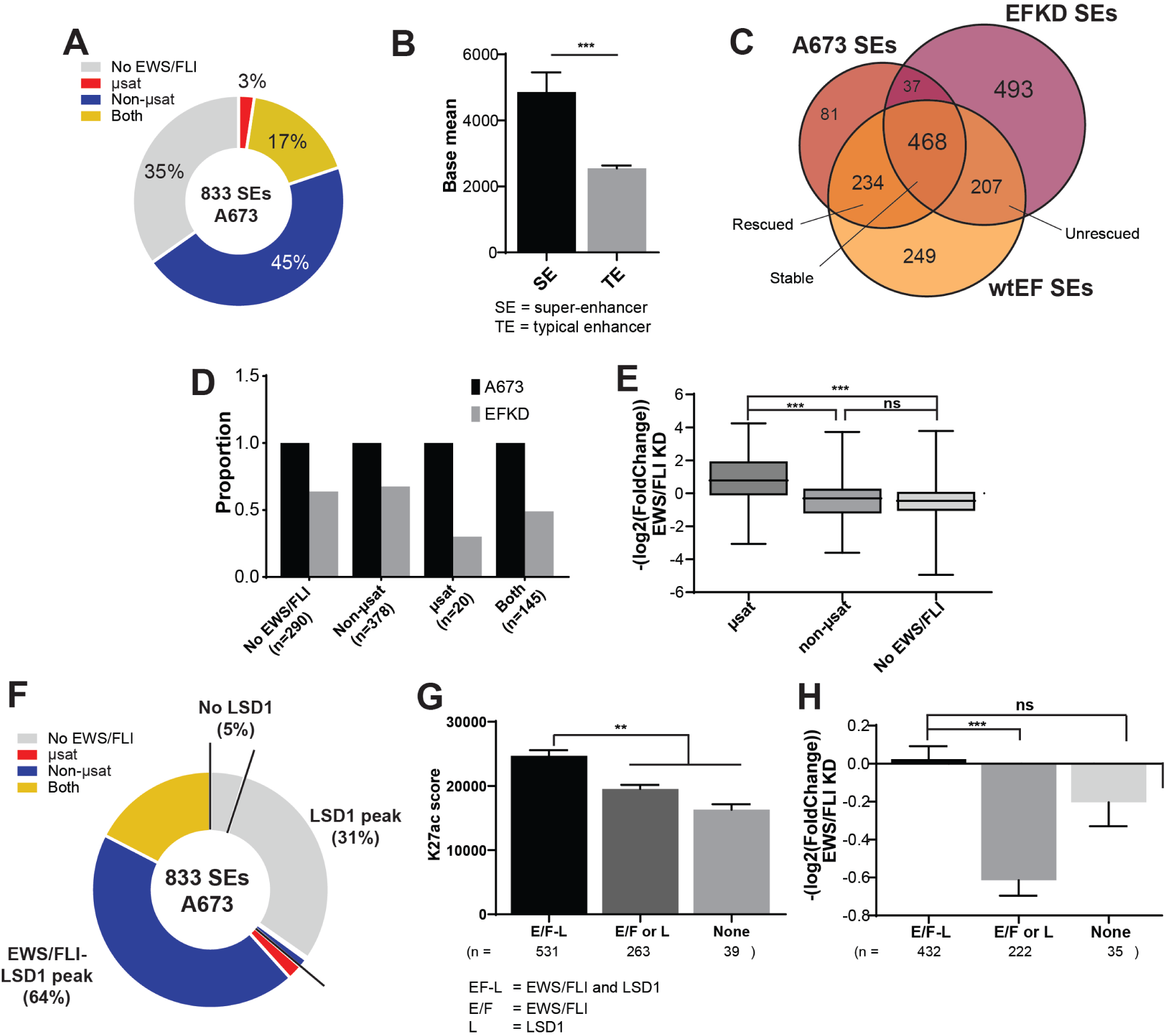
Super-enhancers in A673 cells are associated with both EWS/FLI and LSD1. A) Pie chart distribution of super-enhancers (SEs) in A673 cells by type of overlapped EWS/FLI-bound motif. B) Base mean expression for genes associated with super-(N=615) and typical (N=6958) enhancers in A673 cells. Mean and SD are shown and p-values were determined using an unpaired t-test. ***p<0.001. C) Venn diagram of SEs in A673, EFKD, and wtEF cells as determined by ChIPPeakAnno. D) Proportions of SEs present in A673 and EFKD cells sorted by the type of EWS/FLI-bound motif overlapped by the SE. E) EWS/FLI-mediated differential expression for genes associated with SEs in A673 cells sorted by the type of EWS/FLI-bound motif overlapped by the SE. Mean and SD are shown and p-values were determined using one-way ANOVA with multiple comparison testing (***p<0.001, **p<0.01, *p<0.05.) F) Pie chart distribution of SEs by type of EWS/FLI and LSD1 overlap. G,H) (G) H3K27ac score calculated from the ROSE algorithm and (H) EWS/FLI-mediated differential expression of nearby genes for SEs in A673 cells plotted by type of overlap with EWS/FLI and LSD1. EF-L=EWS/FLI and LSD1 coincident peak, E/F=EWS/FLI only, L=LSD1 only. Mean and SD are shown. N for differential expression and base mean is lower for those K27ac scores because not all genes near SEs were detected by RNA-seq. P-values were determined using one-way ANOVA with multiple comparison testing (***p<0.001, **p<0.01, *p<0.05.) See also Supplementary Figures 9-11 and Supplementary Tables 1-14.

*De novo* establishment of enhancers at GGAA-µsats is unique to Ewing sarcoma, due to the altered binding specificity conferred to the FLI DBD in the fusion.^12,42^ Because of the observation that SEs in Ewing sarcoma cells contained both GGAA-µsats and non-µsats, we next investigated how the type of EWS/FLI binding site (or sites) contained within a SE determined the stability of that SE in the absence of EWS/FLI. Following EWS/FLI-depletion (Supplementary Figure 9B-C), 315 (38%) of all SEs collapsed and 700 new SEs were established in EFKD cells (Figure 3C, Supplementary Figure 9D; Supplementary Tables 3-4). Of the 315 SEs which collapsed 234 (74%) were reconstituted in cells rescued with ectopic EWS/FLI expression (wtEF cells) (Figure 3C). Rescue also resulted in 493 (70%) of the 700 EFKD-specific SEs collapsing (Figure 3C, Supplementary Figure 9E; Supplementary Tables 5-6). SEs which contained any GGAA-µsat were less stable in EFKD cells (46% persist) than SEs either overlapping only a non-µsat EWS/FLI site or containing no EWS/FLI binding (non-µsat only: 67% persist, no EWS/FLI: 64% persist, Figure 3D). This was true for both GGAA-µsat-containing SEs which overlapped a non-µsat (“both”; 49% persist, Figure 3D) and those that did not (“µsat”; 30% persist, Figure 3D). These data suggest GGAA-µsat-associated SEs are more dependent on EWS/FLI-binding and are more likely to collapse when EWS/FLI is depleted compared to non-µsat-associated SEs.

Consistent with these data, genes nearest to SEs containing an EWS/FLI-bound GGAA-µsat were upregulated by EWS/FLI as compared to those near SEs lacking a GGAA-µsat, which were slightly downregulated (Figure 3E). These data suggest that the transcriptional machinery may preferentially accumulate at SE loci bound by EWS/FLI at GGAA-µsats, effectively sequestering these complexes away from other SEs and leading to a reduction of transcription levels where SEs do not contain a GGAA-µsat.

### LSD1 enhances the establishment of super-enhancers by EWS/FLI

Having observed that many Ewing sarcoma SEs also contain non-µsat sites, we next asked whether LSD1 was also present in these SEs, and whether the LSD1 harbored in SEs might be colocalized with, or bind independently from, EWS/FLI. In other non-Ewing sarcoma contexts, LSD1 is enriched at SEs^43^, though its function is unclear. LSD1 is also implicated in genome-wide maintenance of primed enhancers.^44^ We found 95% of A673 SEs overlapped an LSD1 peak. There were 64% of SEs overlapping a locus with colocalized EWS/FLI and LSD1, while 31% of SEs were overlapping an LSD1 peak without any colocalized EWS/FLI (Figure 3F, Supplementary Table 2).

To determine the functional relationship between LSD1, EWS/FLI, and SEs in A673 cells, we analyzed SEs based on their EWS/FLI and LSD1 binding status. SEs possessing an EWS/FLI-LSD1 coincident peak had significantly higher H3K27ac scores than SEs containing either non-overlapping EWS/FLI and/or LSD1 peaks (p<0.01), or neither EWS/FLI nor LSD1 (p<0.01; Figure 3G). While there was no significant difference in base expression of the nearest gene (Supplementary Figure 9F), those SEs which lacked EWS/FLI-LSD1 co-peaks showed decreased gene expression in the presence of EWS/FLI (Figure 3H). That the highest levels of H3K27ac were seen at SEs where EWS/FLI and LSD1 are colocalized, and that genes near SEs which lack this colocalization tend to be downregulated by EWS/FLI, suggests that the cooperation between EWS/FLI and LSD1 promotes deposition of H3K27ac and may lead to preferential accumulation of transcriptional machinery.

Having focused primarily on SEs in A673s cells, we next asked whether these relationships between EWS/FLI, LSD1, and histone H3K27ac deposition was a common feature across Ewing sarcoma cell lines. We found that a majority of SEs overlapped non-µsat-bound EWS/FLI, either with or without overlap of a GGAA-µsat in A673, EWS-502, and SK-N-MC cells (Supplementary Figure 10A-C). TC71 cells had fewer SEs overlapping EWS/FLI, but most of those TC71 SEs with EWS/FLI binding were also overlapping a non-µsat. (Supplementary Figure 10D). A significant portion of SEs overlapping EWS/FLI also overlapped with LSD1 binding (Supplementary Figure 10E-H) and H3K27ac deposition was highest at those SEs with EWS/FLI-LSD1 colocalization (Supplementary Figure 10I-L) across all cell lines.

Other studies have suggested LSD1 acts genome-wide to maintain active and primed enhancers, and it is proposed that LSD1 does this by functioning as a repressor and preventing over-activation.^44^ In order to clarify whether LSD1 at EWS/FLI-activated enhancers was simply part of this repressive maintenance function, or instead whether LSD1 promoted enhancer activity, we used GSEA to ask how LSD1 regulated genes near SEs with colocalized EWS/FLI and LSD1. We found that these genes were functionally associated with LSD1-mediated gene activation across all cell lines (Supplementary Figure 11A-D), suggesting that LSD1 is not functioning as a repressor at the enhancers associated with these genes. Together, these results show LSD1 colocalized at EWS/FLI-bound non-µsat loci correlates with increased H3K27ac deposition. This occurs regardless of whether the SE also overlaps an EWS/FLI-bound GGAA-µsat. A model for this will be more fully described in the Discussion section below.

### EWS/FLI causes dynamic reorganization of LSD1 genome-wide

Functional association of EWS/FLI with LSD1 could occur through 1) active redistribution of LSD1 caused by EWS/FLI or 2) binding of EWS/FLI at loci preloaded by LSD1 in a precursor cell. To determine which of these mechanisms operates in Ewing sarcoma, we evaluated LSD1 occupancy in either parental A673 cells or EFKD cells. We additionally included EFKD cells rescued with ectopic expression of EWS/FLI, wtEF cells. Panels show specific examples at *LMO2* and *SERPINE1* where EWS/FLI depletion drives reversible changes in LSD1 binding in Figure 4A-B and Supplementary Figure 12. Globally, LSD1 was bound at 40262 loci in A673 cells, 33085 loci in EFKD cells, and 39659 loci in wtEF cells (Figure 4C). We observed 16698 LSD1 peaks present in A673 cells that collapse in EFKD cells, 9197 of which are rescued in wtEF cells (Figure 4C). Of the 10151 loci which gain LSD1 peaks following EWS/FLI depletion, 7459 loci lose LSD1 binding upon rescue with ectopic EWS/FLI expression. Notably, while we initially observed 21950 LSD1-bound loci were “stable” across the tested conditions, a closer inspection revealed more dynamism within these stable peaks than we had appreciated. Of these “stable” peaks, 5687 show increased LSD1 binding in A673 as compared to EFKD, and 9271 show greater binding in EFKD cells as compared to A673, further supporting that EWS/FLI expression results in genome-wide reorganization of LSD1 (Figure 4D).

**Figure 4.**
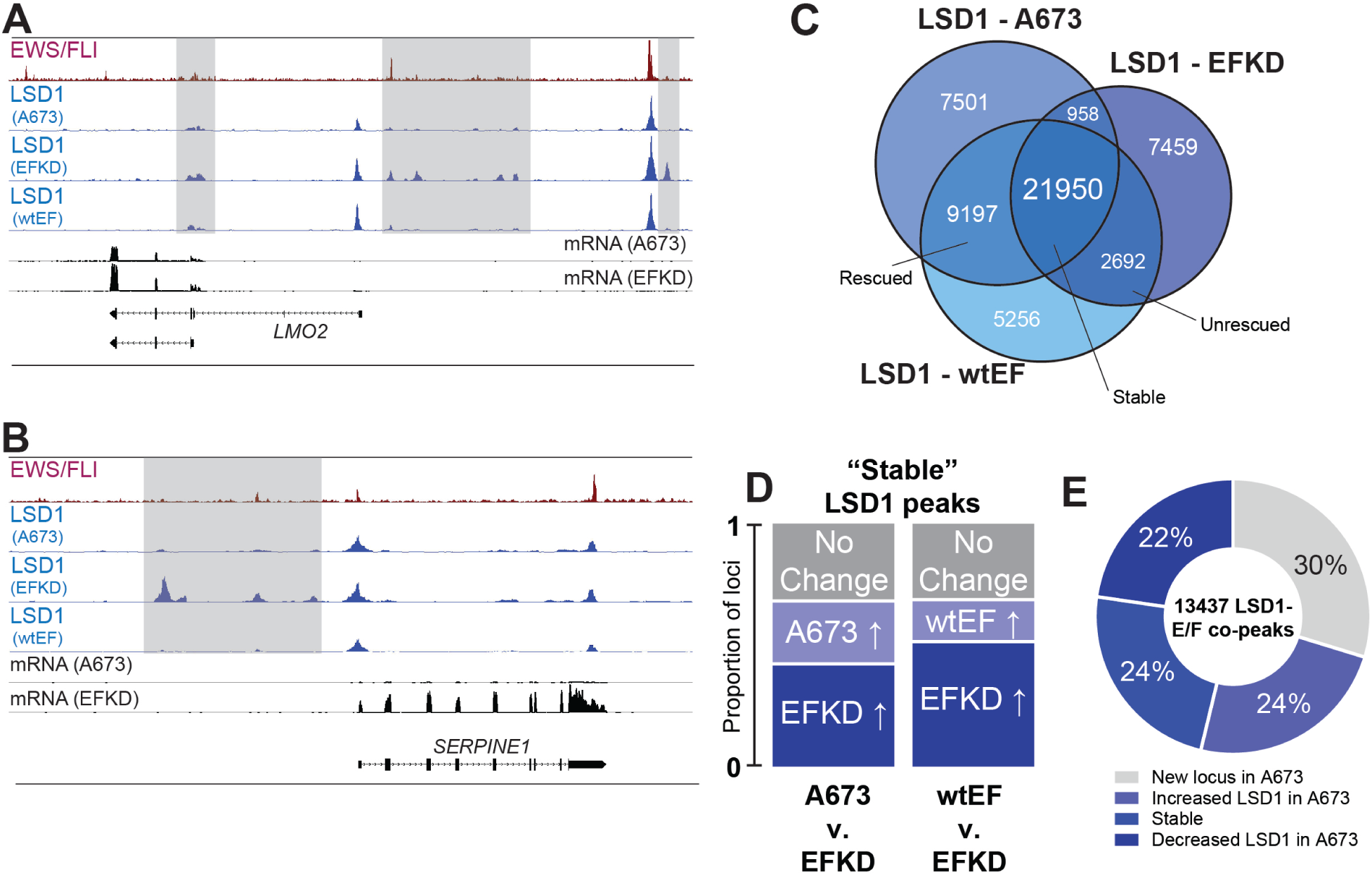
EWS/FLI alters the genome-wide occupancy of LSD1. A,B) IGB tracks showing EWS/FLI and LSD1 near (A) *LMO2* and (B) *SERPINE1*. Tracks show LSD1 in A673, EFKD, and wtEF cells and mRNA in the A673 and EFKD conditions. C) Venn diagram of LSD1 peaks in A673, EFKD, and wtEF cells as determined by ChIPPeakAnno. D) Bar charts showing the dynamics of relative proportions of “stable” LSD1 peaks (detected in A673, EFKD and wtEF). E) Pie chart distribution showing proportion of EWS/FLI-LSD1 coincident peaks with LSD1 binding dynamics as compared to LSD1 localization in EFKD cells. See also Supplementary Figure 12.

Considering the widespread redistribution of LSD1, we next asked whether LSD1 colocalizes with EWS/FLI at new sites in the genome, or if EWS/FLI instead binds at loci which already possess LSD1. Venn diagram analysis showed most LSD1 peaks present in A673 cells were also present in EFKD cells (Figure 4C). Of the EWS/FLI-LSD1 colocalized peaks, 54% show increased LSD1 binding in A673 cells, with LSD1 binding at a new locus with EWS/FLI 30% of the time, while another 24% show increased LSD1 binding with EWS/FLI expression (Figure 4E). These data suggest both that LSD1 is recruited to new sites and that EWS/FLI binds at sites already bound by LSD1. At these latter sites, we speculate that LSD1 may interact with other ETS factors when EWS/FLI is absent and that EWS/FLI may displace these ETS factors, as has been previously suggested,^8^ and hijack LSD1 activity.

### LSD1 binds at activating “super-clusters”

We were struck by the visual clustering of LSD1 peaks in cells with depleted EWS/FLI expression, as shown in Figure 4A and 4B. Because clustering of chromatin regulatory proteins, including LSD1, is reported at SEs, we investigated the relationship between LSD1 “super-clusters” (SCs) and SEs in EFKD cells. The ROSE algorithm identified 970, 1287, and 1325 LSD1 SCs in A673, EFKD, and wtEF cells, respectively (Figure 5A, Supplementary Tables 15-20). These are regions with the highest levels of LSD1 binding throughout the genome, as defined by a function of rank and LSD1 signal (Figure 5B-C, Supplementary Figure 13A). Reflecting global LSD1 binding, we observed EWS/FLI-driven dynamism in the genome-wide distribution of LSD1 SCs. There were 426 LSD1 SC’s present in A673 cells that collapse in EFKD cells, 269 of which are rescued in wtEF cells (Figure 5A). Of the 753 loci which gain LSD1 clusters following EWS/FLI depletion, 498 loci lose LSD1 binding upon rescue. LSD1 clusters are stable regardless of EWS/FLI status at 486 loci.

**Figure 5.**
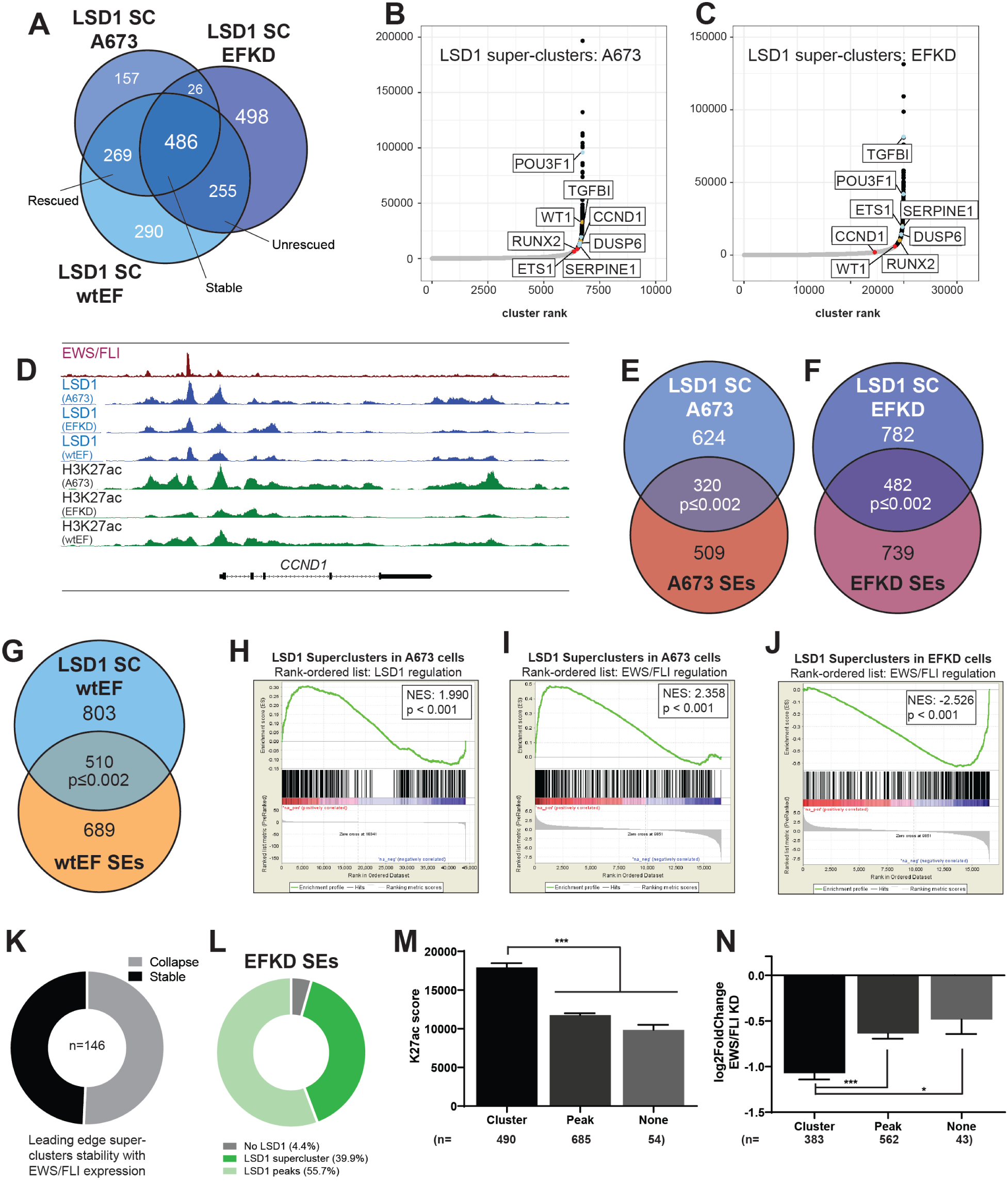
LSD1 binds in super-clusters that are disrupted by EWS/FLI. A) Venn diagram of LSD1 SCs in A673, EFKD, and wtEF cells B,C) Plotted output of the ROSE analysis for LSD1 superclusters (SCs) in (B) A673 and (C) EFKD cells. D) IGB tracks showing coincidence of EWS/FLI, LSD1, and H3K27ac in a SE and LSD1 SC near *CCND1*. Tracks show LSD1 and H3K27ac in A673, EFKD, and wtEF conditions. E-G) Venn diagrams of SEs and LSD1 SCs in (E) A673 cells, (F) EFKD cells, and (G) wtEF cells. Overlaps and p-values were determined by ChIPPeakAnno. H-J) GSEA results using genes near (H,I) LSD1 SCs in A673 cells (N=427) or (J) EFKD cells (N=500) as the test set and either LSD1 gene regulation in A673 cells (H) or EWS/FLI gene regulation (I,J) as the rank-ordered dataset. NES=normalized enrichment score. |NES|>1.5 is significant. K) Pie chart distribution showing the number of leading edge LSD1 SCs (from J) that collapse in A673 cells. L) Pie chart distribution showing the overlap of SEs in EFKD cells with different types of LSD1-binding. M,N) (M) H3K27ac score calculated from the ROSE algorithm and (N) EWS/FLI-mediated differential expression of genes near SEs in EFKD cells plotted by type of overlap with LSD1. Mean and SD are shown. N for (N) is lower than (M) because not all genes near SEs were detected by RNA-seq. p-values were determined using one-way ANOVA with multiple comparison testing (***p<0.001, **p<0.01, *p< 0.05.) See also Supplementary Figures 13-14 and Supplementary Tables 2, 4, 6, and 15-20.

Due to notable overlaps between LSD1 SCs and SEs at individual loci, such as those shown at *CCND1* (Figure 5D), *DUSP6* (Supplementary Figure 13B), *ETS1* (Supplementary Figure 13C), and *TGFBI* (Supplementary Figure 13D), we initially considered whether LSD1 SCs simply represented SEs. However, only 320 A673 SCs (33%) and 482 EFKD SCs (37%), and 510 wtEF SCs (38%) overlapped with SEs in their respective cells (Figure 5E-G), instead suggesting a heretofore unappreciated chromatin-associated LSD1-organizational structure. GSEA revealed that LSD1 SCs were associated with both LSD1-mediated gene activation (NES=1.990, p<0.001; Figure 5H) and EWS/FLI-mediated gene activation (NES=2.358, p<0.001; Figure 5I) in A673 cells, consistent with prior observations that LSD1 plays a role in EWS/FLI-mediated activation. In contrast, in EFKD cells, LSD1 SCs were strongly associated with genes that are repressed by EWS/FLI and thus become activated in the knockdown condition (NES=-2.526, p<0.001; Figure 5J), again supporting a role for LSD1 in gene activation, even in the absence of EWS/FLI. Most of the genes in the leading edge of this latter GSEA have SCs that collapse with wildtype levels of EWS/FLI (Figure 5K). We speculate that LSD1 SCs are associated with gene activation in a Ewing sarcoma precursor cell, and that during the process of Ewing sarcoma development these activating LSD1 SCs collapse and expression of nearby genes is downregulated. In Ewing sarcoma cells new LSD1 SCs are formed.

Although the overlap of LSD1 SCs with SEs in EFKD cells was partial, we found an overwhelming majority (95.6%) of SEs in these cells overlapped at least one LSD1 peak (Figure 5L, Supplementary table 19). This was similar to our prior observations of SEs in A673 cells (Figure 3F). 39.9% of super-enhancers overlapped an LSD1 SC, while 55.7% overlapped a “monopeak” (LSD1 bound in an individual peak, not as part of a cluster). A similar distribution was seen for SEs in A673 and wtEF cells (Supplementary Figure 14A-B). In all conditions, those SEs overlapping an LSD1 SC had greater H3K27ac scores than those with only a monopeak, or no LSD1 (Figure 5M, Supplementary Figure 14C-D), suggesting a functional role for the factors that recruit LSD1 in promoting the establishment of enhancers. No significant difference was observed in basal expression of SE-associated genes based on LSD1 binding status (Supplementary Figure 14E-G), but genes near EFKD SEs containing an LSD1 SC showed greater downregulation with EWS/FLI expression than genes near SEs without an LSD1 SC (Figure 5N). In cells expressing EWS/FLI, EWS/FLI-mediated regulation of SE-associated genes showed no such dependency on LSD1 configuration within the SE (Supplementary Figure 14H-I). Interestingly, of the 251 SEs that both 1) are unique to EFKD and 2) overlap an LSD1 SC, 196 (78%) have SCs that are also unique to EFKD cells (Supplementary Figure 14J), indicating a concurrent collapse of both the LSD1 SC and the SE upon EWS/FLI expression. Taken together, these data support a second novel model for EWS/FLI-mediated repression via aberrant enhancer regulation: EWS/FLI-induced LSD1 SC collapse prevents priming and maintenance of enhancers active in the Ewing sarcoma precursor cell. Moreover, once EWS/FLI is introduced to the cell, the primacy of EWS/FLI-mediated transcriptional regulation overtakes that of LSD1-SCs in the determination of gene expression.

## DISCUSSION

The close phenotypic overlap between LSD1 inhibition (with SP2509) and EWS/FLI depletion in A673 (or EWS/ERG depletion in TTC-466 cells) suggested that LSD1 is closely linked to the genome-wide activity of oncogenic fusions in Ewing sarcoma.^17^ Prior studies suggested that LSD1 is part of a NuRD-LSD1 complex hijacked by EWS/FLI to repress tumor suppressors, but how LSD1 was involved in EWS/FLI-mediated gene activation was unclear.^9^ To understand this relationship, we used genomic approaches to probe LSD1 distribution and function in four Ewing sarcoma cell lines, and a model of the Ewing sarcoma precursor cell with diminished EWS/FLI expression, EFKD, complemented with rescue using ectopic EWS/FLI expression, wtEF. Though EFKD cells are an imperfect precursor model, we believe they are both conceptually and technically useful in that they are a system which tolerates EWS/FLI (re-)introduction. Importantly, following EWS/FLI depletion, they continue to proliferate^10^, enabling the requisite large numbers of cells needed for chromatin-level analyses.

We found that LSD1 is broadly important for gene activation, functioning at enhancers in both Ewing sarcoma and precursor cells, and that EWS/FLI drives dynamic genome-wide reorganization of LSD1. Functional interaction between EWS/FLI and LSD1, particularly at non-µsat sites, is critical to restructure the enhancer landscape in Ewing sarcoma cells as modeled in Figure 6. Here, we build on previous studies that show *de novo* enhancer formation at EWS/FLI-bound GGAA-µsats and found that these enhancers almost always also involve an EWS/FLI-bound HA site (Figure 6A Panel i). LSD1 is frequently recruited to these collaborating loci and the presence of LSD1 augments enhancer formation, resulting in increased H3K27ac deposition. Panel ii depicts enhancers which are solely driven by HA sites. In precursor cells, these are likely bound by other ETS transcription factors and LSD1. In Ewing sarcoma cells, it is probable that EWS/FLI hijacks these sites through displacement of the endogenous ETS factor while retaining LSD1 binding.

**Figure 6.**
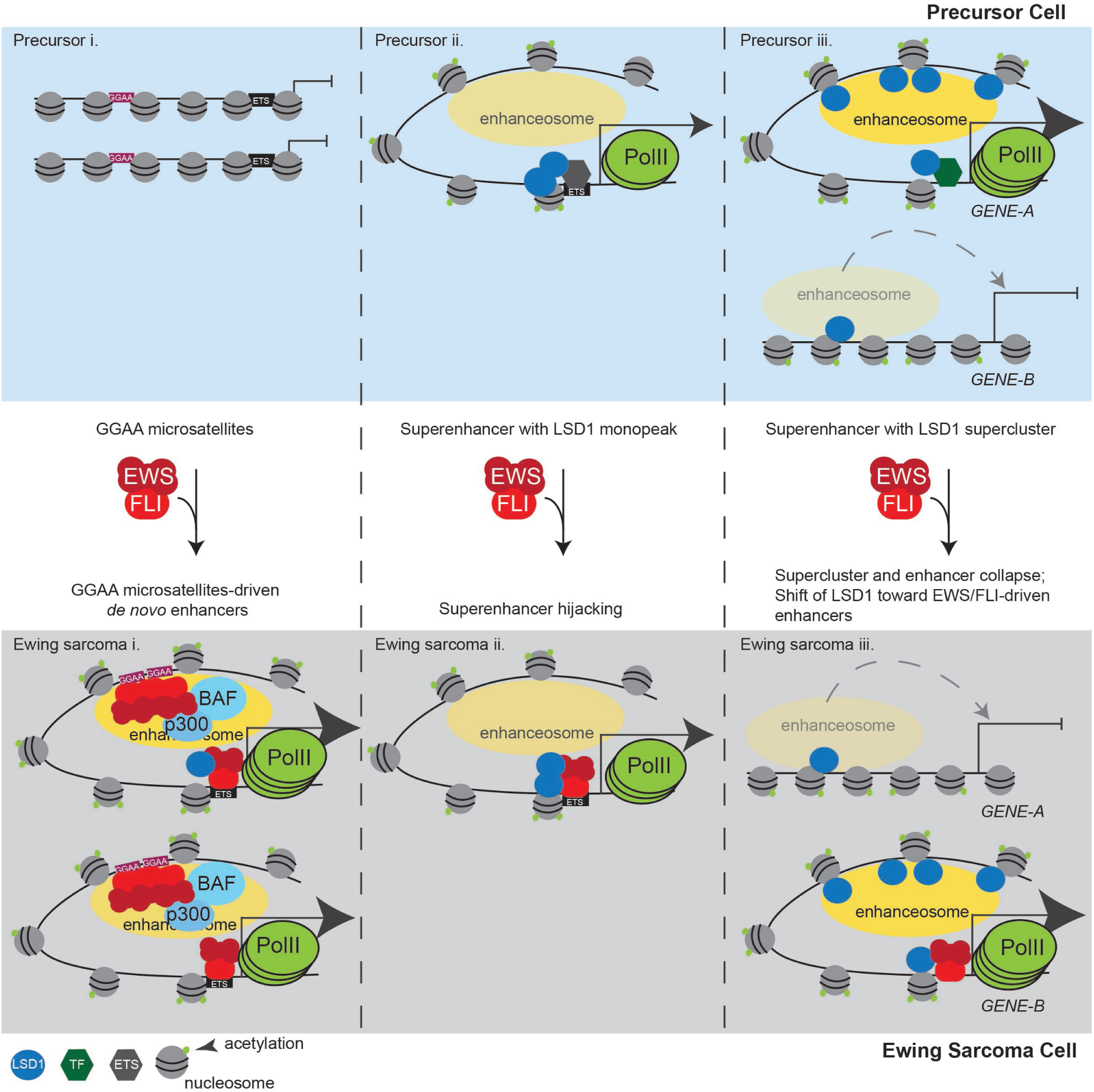
LSD1 is tightly linked to the shifting enhancer landscape in Ewing sarcoma. A) Model figure showing how EWS/FLI remodels the enhancer landscape and the role of LSD1 in this remodeling. The top panels depict enhancer states found in a precursor cell and the bottom panels represent a Ewing sarcoma cell. Panel (i) shows chromatin remodeling which results in de novo enhancer formation at GGAA-µsats. Panel (ii) shows chromatin remodeling which occurs at enhancers bound by LSD1 with another ETS family member in precursor cells. These enhancers are hijacked by EWS/FLI. Panel (iii) shows supercluster and enhancer collapse which occurs at enhancers with LSD1 superclusters in precursor cells with establishment of an LSD1-decorated supercluster driven by EWS/FLI. The number of PolII molecules by any gene correlates to the level of transcription from those genes.

The dynamic reorganization of LSD1 SCs is shown in Panel iii. In precursor cells, LSD1 SCs promote nearby enhancer formation and gene activation. Expression of EWS/FLI disrupts these loci, causing collapse of LSD1 SCs and the associated super-enhancers, leading to downregulation of nearby genes. EWS/FLI thus engages distinct mechanisms to alter the function of LSD1-containing complexes: 1) through direct recruitment of the NuRD-LSD1 complex previously described^9^ and 2) through reorganization of LSD1 and LSD1 SCs.

This model is compelling because it enhances our understanding of aberrant epigenomic regulation driven by EWS/FLI. Our results suggest that EWS/FLI-bound GGAA-µsats may depend upon another EWS/FLI binding event at a non-µsat to target the enhancer activity. Recruitment of LSD1 to these non-µsat sites further augments EWS/FLI-mediated enhancer formation, and this occurs even in the absence of GGAA-µsats. These findings unite important observations regarding the involvement of both GGAA-µsats and LSD1 in EWS/FLI-mediated gene activation. We further identified two novel mechanisms for gene downregulation by EWS/FLI, and both are intricately linked to altered enhancer function. First, there exist some SEs which show decreased transcriptional activity with EWS/FLI expression. These SEs frequently do not overlap either colocalized EWS/FLI-LSD1 or a GGAA-µsat, suggesting that transcriptional machinery preferentially accumulates at SEs where EWS/FLI both binds a GGAA-µsat and is colocalized with LSD1, while transcriptional machinery is depleted at other SEs lacking EWS/FLI binding. Second, EWS/FLI-induced collapse of LSD1 SCs leads to decreased enhancer priming. Despite the strong transcriptional activation capacity of the EWS domain, expression of EWS/FLI results in a greater number of genes repressed than activated, and these two mechanisms likely contribute to this process.

LSD1 is important for enhancer decommissioning during differentiation^23^ and LSD1 constructs fused to transcription activator-like effector (TALE-LSD1) or enzymatically dead Cas9 (dCas9-LSD1) show that LSD1 silences enhancers and promoters when targeted to specific genomic loci.^45,46^ More recent studies highlight a role for LSD1 involvement in enhancer silencing by lineage-specific transcription factors like GFI1 in acute myeloid leukemia^25^ and medulloblastoma^47^, or BCL6 in diffuse large B-cell lymphoma.^24^ In these cases, inhibition of LSD1 with derivatives of tranylcypromine restores enhancer function and disrupts oncogenic gene regulation. However, we observed LSD1 to be largely associated with gene activation in Ewing sarcoma. Indeed, knockdown of LSD1 results in the downregulation of activated genes nearby,^34^ indicating that LSD1 is not functioning to suppress over-activation, but is instead critically important to maintain gene activation. How EWS/FLI enforces an activating role for LSD1 is unknown, but the activity observed is similar to LSD1 activity in prostate cancer. In prostate cancer LSD1 activates oncogenic gene transcription independently from its enzymatic function.^48,49^ Interestingly, both Ewing sarcoma and prostate cancer show sensitivity to reversible LSD1 inhibition with SP2509, but not other classes of irreversible LSD1 inhibitors related to tranylcypromine.^34,49^ This suggests that different functions of LSD1 may be differentially targeted by different classes of LSD1 inhibitors. The specific mechanistic role LSD1 is playing here, whether non-enzymatic or through demethylation of targets other than H3K4, is not yet known and remains an important area of future study.

In conclusion, EWS/FLI interacts with LSD1 to mediate genome-wide epigenetic and transcriptional changes in Ewing sarcoma. EWS/FLI induces a dynamic reorganization of LSD1 that acts in concert with EWS/FLI activity at GGAA-µsats to reshape the enhancer landscape. The involvement of widespread localization of LSD1 at EWS/FLI-bound non-µsats suggests that EWS/FLI-mediated chromatin regulation in Ewing sarcoma requires widespread activity at loci beyond GGAA-µsats. The mechanisms which drive this non-µsat-mediated regulation are poorly understood and represent critical facets of EWS/FLI function to explore. We also show that LSD1 binds chromatin in a “clustered” configuration. While a similar binding pattern has been observed for LSD1 enriched at SEs, we found an imperfect overlap between LSD1 clusters and SEs. This study suggests that understanding how these clusters form and function, and how perturbations occur in disease, could provide clues on how to better target LSD1 function in Ewing sarcoma patients, as well as in other malignancies.

## MATERIALS AND METHODS

### Key Resources

Key resources required for this protocol are listed in Supplementary Table 21.

### Cell Lines

All cell lines included are tested for mycoplasma annually and sent for STR profiling every two years. All cell lines recently tested negative for mycoplasma and were most recently authenticated by STR profiling in 2018. We should note that we have used the SK-N-MC and A673 lines. These were previously misidentified as neuroblastoma and rhabdomyosarcoma lines, respectively, but actually contain the EWS/FLI fusion and are Ewing sarcoma cell lines.

All Ewing sarcoma cells were cultured at 37°C, 5% CO_2_. A673 and SK-N-MC cells were cultured in DMEM (Corning Cellgro 10-013-CV) containing 10% fetal bovine serum (FBS, Gibco 16000-044), penicillin/streptomycin/glutamine (PSQ, Gibco 10378-016), and sodium pyruvate (Gibco 11360-070). EWS-502 and TC71 cells were cultured in RPMI (Corning Cellgro 15-040-CV) containing 10% FBS for TC71 cells and 15% FBS for EWS-502 cells, as well as P/S/Q. A673 cells are derived from the tumor of a 14-year old Japanese female, contain a type I EWS/FLI fusion, have mutant *TP53* (Q119fs) and wildtype *STAG2*. EWS-502 cells are derived from a Ewing sarcoma patient of unspecified sex and age, and have mutant *TP53* (C135F) and *STAG2* loss. SK-N-MC cells are derived from the tumor of a 12-year old female, have truncated *TP53*, and wildtype *STAG2*. TC71 cells are derived from the tumor of a 22-year old male, have mutated *TP53* (R213*), and have wildtype *STAG2*.

HEK293-EBNA cells were grown at 37°C, 5% CO_2_ in DMEM supplemented with 10% FBS, penicillin/streptomycin/glutamine, and 0.3 mg/mL geneticin (Gibco 10131-027). These cells are derived from the kidney of a healthy aborted fetus, presumed female. Cells were originally transformed by culturing with sheared adenovirus 5.

### Retrovirus Production

To generate retroviruses of the previously reported constructs for iLuc and iEF-2 shRNAs, as well as cDNA for 3XFLAG-Δ22 and 3XFLAG-EWS/FLI^10^, HEK293-EBNA cells were co-transfected with retroviral expression plasmids, vesicular stomatitis virus G glycoprotein (VSV-G) and gag/pol packaging plasmids using Mirus Bio TransIT-LT1. Following 48 hours virus-containing supernatant was collected and filtered. Retrovirally infected A673 cells were selected in 2 µg/mL puromycin (Sigma P8833) for a minimum of 72 hours. For rescued cells, infection occurred after 72 hours of puro selection and cells were double selected for 7 additional days in puro with 100 µg/mL hygromycin B.

### Immunoblotting

For validation of protein knockdown, samples were run on 4-15% Mini-PROTEAN TGX precast gels (BioRad) using 90V for 15 minutes and 120 V for 50 minutes. Proteins were blotted to nitrocellulose membranes using semi-dry transfer with the Bjerrum Schaffer-Nielsen buffer at 15 V for 60 minutes. Membranes were blocked at 4°C overnight in Odyssey Blocking Buffer PBS (LI-COR), and incubated with primary antibody overnight at 4°C. Primary antibodies used for immunoblotting were: anti-FLI (Abcam ab15289), anti-H3 (Cell Signaling Technology #4499 - D1H2), anti-Lamin B1 (Abcam ab16048), and anti-FLAG M2 (Sigma F3165). For validation of protein depletion with knockdown, FLI, total H3, Lamin, and FLAG blots were incubated with IRDye secondary antibodies (LI-COR) and developed on the Odyssey.

### Chromatin Immunoprecipitation and Sequencing (ChIP-seq)

For performing chromatin immunoprecipitation, A673 cells were seeded in 15-cm dishes. Cells were removed by scraping in plain media and pelleted at 1200 rpm for 5 min. Pellets were resuspended in room temperature cell lysis buffer (20 mM HEPES-KOH, pH 8.0; 1 mM EDTA; 0.5 mM EGTA; 140 mM NaCl; 10% glycerol; 0.5% NP-40; 0.25% Triton X-100; protease inhibitor) and incubated on ice for 5 min before nuclei were pelleted at 1200 rpm for 5 min. Nuclei were washed once in 1 mL MNase digestion buffer (20 mM Tris-Cl. pH 7.4; 5 mM MgCl_2_; 1 mM CaCl_2_; 0.1% Triton X-100, protease inhibitor) and resuspended in MNase digestion buffer. 50 U (10 µL of 5 U/µL) MNase were added and nuclei were incubated at 37°C for 30 minutes with gentle inversion every 10 minutes. Digestion was stopped by the addition of chilled 100 µL 0.5M EDTA followed by a 5-minute incubation on ice. Nuclei were cleared with a spin at 13000 rpm for 10 minutes and the resulting supernatant (chromatin) was transferred to a new tube. Chromatin concentration was quantified by Nanodrop and 100 µg of chromatin was used for each immunoprecipitation, diluted to equal volumes in dilution buffer (20 mM Tris-Cl, pH 7.4; 2 mM EDTA; 50 mM NaCl; 0.25% Triton X-100; 20 mg/mL BSA; protease inhibitor) with 50 µL of 0.5M EDTA. Chromatin was added to antibody coated magnetic Dynabeads for 16-20 hours at 4°C. Antibodies used for ChIP were: anti-H3K4me1 (ab8895), anti-H3K4me2 (ab32356), and anti-H3K4me3 (ab8580). Beads were washed with ChIP wash buffers containing 0.1% SDS and 150 mM NaCl, 150 mM NaCl, and 250 mM LiCl. ChIP DNA was eluted with fresh elution buffer, RNAse and Proteinase K treated and decrosslinked overnight at 65°C. DNA was then purified with phenol-chloroform using the Qiagen MaXtract extraction kit as per manufacturer’s instructions. DNA was quantified by Qubit and validated for enrichment by qPCR using specific primers. Following validation, libraries were generated for sequencing using the NEBnext kit according to manufacturer’s instructions and submitted for deep sequencing on the Illumina HiSeq 4000 platform (Nationwide Children’s Hospital Institute for Genomic Medicine). Native histone ChIPs were performed three times, with a non-specific IgG negative control.

### Cleaveage Under Targets and Release Using Nuclease (CUT&RUN) and Cleavage Under Targets and Tagmentation (CUT&Tag)

#### Cell Preparation

CUT&RUN and CUT&Tag were performed as described^53,54^ with slight modifications. BioMag® Plus Concanavalin A-coated magnetic beads (Bangs Laboratories, BP531; 10 µl beads per condition) were washed twice with Binding buffer (20 mM HEPES-KOH pH 7.9, 10 mM KCl, 1 mM CaCl_2_, 1 mM MnCl_2_) in preparation for CUT&RUN/CUT&Tag. 500,000 cells (CUT&RUN) and 250,000 cells (CUT&Tag) per condition were washed twice with Wash Buffer (20 mM HEPES-NaOH pH 7.5, 150 mM NaCl, 0.5 mM Spermidine, Protease Inhibitor) and rotated with prepared beads for 10 minutes at room temperature. The supernatant was cleared and removed using a magnet stand. The beads were resuspended in 100 µL Antibody Buffer (20 mM HEPES-NaOH pH 7.5, 150 mM NaCl, 0.5 mM Spermidine, 0.02% Digitonin [CUT&RUN] or 0.05% digitonin [CUT&Tag], 2 mM EDTA, Protease Inhibitor) and antibodies (FLI 7.3 mouse, Santa Cruz; H3K27ac rabbit, Abcam ab4729; Rabbit anti-mouse IgG, Abcam ab46540; LSD1 rabbit, Abcam ab17721) were added at a dilution of 1:100. Samples were rotated overnight at 4°C. The samples were cleared on a magnet stand and beads were washed with Dig-wash buffer (20 mM HEPES-NaOH pH 7.5, 150 mM NaCl, 0.5 mM Spermidine, 0.02% Digitonin [CUT&RUN] or 0.05% digitonin [CUT&Tag]).

#### CUT&RUN (FLI, LSD1, H3K27ac, and Rb IgG)

Beads that were incubated with FLI 7.3 mouse antibody were resuspended in 100 µL Dig-wash buffer and incubated with rabbit anti-mouse secondary antibody (Abcam, ab46540) at a dilution of 1:100 on a rotator for 1 hour at 4°C. All other samples didn’t require a secondary antibody step. After another wash with Dig-wash buffer, beads were resuspended in 100 µL Dig-wash buffer and Protein A-MNase fusion protein (generously provided by the Henikoff lab) was added to a final concentration of 700 ng/mL. Samples were rotated for 1 hour at 4°C. After 2 washes with Dig-wash buffer, beads were resuspended in 100 µL Dig-wash buffer and placed in ice water to equilibrate to 0°C. CaCl_2_ was added to a final concentration of 2 mM under gentle vortexing and samples were incubated for 45 minutes (H3K27ac) or 2 hours (FLI, LSD1) at 0°C. Reactions were stopped by adding 100 µl 2XSTOP buffer (340 mM NaCl, 20 mM EDTA, 4 mM EGTA, 0.02% Digitonin, 0.05 mg/mL RNase A, 0.05 mg/mL Glycogen containing 2 pg/mL heterologous Yeast Spike-in DNA) and incubated at 37°C for 10 minutes to release the CUT&RUN fragments. Beads were pelleted by centrifugation at 16,000 x g and 4°C for 5 minutes and supernatants containing CUT&RUN fragments were transferred to new tubes. SDS was added to a final concentration of 0.1% and Proteinase K to a final concentration of 0.25 µg/µL followed by an incubation at 70°C for 10 minutes. DNA from all supernatants was purified using Phenol/Chloroform extraction and ethanol precipitation.

The library prep was performed using the KAPA Hyper Prep Kit (KAPA Biosystems, KK8502) in combination with the KAPA Dual-Indexed Adapter Kit (KAPA Biosystems, #KK8722) with several modifications. 50 µL of CUT&RUN sample were used for the End repair and A-tailing step. A 1.5 µM adapter stock was used for the adapter ligation reaction with a 20-minute incubation step at 20°C. After the recommended post-ligation cleanup, the DNA was eluted in 53 µL elution buffer and a second cleanup was performed using 50 µL of eluted DNA and 65 µL Agencourt AMPure XP magnetic beads (Beckman Coulter, A63880). The DNA was eluted with 25 µL elution buffer and 20 µL were used for the library amplification. To favor small fragments, the amplification was performed using a combined Annealing/Extension step at 60°C for 10 seconds and 13 cycles. 50 µL of the amplified library and 57.5 µL AMPure beads (1.15X) were used for the first post-amplification cleanup. After eluting the DNA with 53 µL, a second post-amplification cleanup step was performed using 50 µL eluted DNA and 62.5 µL AMPure beads (1.25X). The final library was eluted from the beads with 35 µL elution buffer. 2 x 150 bp paired-end sequencing was performed using the Illumina HiSeq4000 system (Nationwide Children’s Hospital Institute for Genomic Medicine). Two independent replicates were performed for each sample, with one replicate consisting of cells prepped from viral infection to sequencing.

#### CUT&Tag (LSD1, H3K27ac, and Rb IgG)

Beads were resuspended in 100 µL Dig-wash buffer and incubated with guinea pig anti-rabbit IgG (Antibodies-Online ABIN101961) at a dilution of 1:100 on a rotator for 1 hour at 4°C. After 3 washes with Dig-wash buffer, beads were resuspended in 100 µL Dig-300 buffer (20 mM HEPES-NaOH pH 7.5, 300 mM NaCl, 0.5 mM Spermidine, 0.01% Digitonin) with a 1:250 dilution of Protein A-Tn5 transposase fusion protein (generously provided by the Henikoff lab). Samples were rotated for 1 hour at room temperature. After 3 washes with Dig-300 buffer, beads were resuspended in 300 µL Tagmentation buffer (Dig-300 buffer with 10 mM MgCl_2_) and incubated for 1 hour at 37°C. Tagmentation was stopped by adding 10 µL 0.5M EDTA, 3 µL 10% SDS, and 2.5 µL 20 mg/mL Proteinase K to each sample, vortexing 5 s, and incubating for 1 hour at 50°C. DNA from samples was directly extracted using phenol-chloroform with ethanol precipitation. Once ethanol-precipitated pellets were dry, pellets were resuspended in 30 µL 10 mM Tris-Cl, pH 8 with 1 mM EDTA and 1/400 RNase A and incubated at 37°C for 10 min.

Libraries were amplified using primers as previously described.^55^ 21 µL of DNA, and 2 µL each of primer (10 µM) were added to 25 µL of NEBNext HiFi 2X PCR master mix and libraries were amplified as follows: 72°C for 5 min, 98°C for 30 s, 15 cycles of 98°C for 10 s and 63°C for 10 s, 72°C for 1 min. After the amplification a cleanup was performed by adding 55 µL Agencourt AMPure XP magnetic beads (Beckman Coulter, A63880) to the PCR reactions, incubating 15 minutes, and washing twice with 400 µL 80% ethanol, and eluting DNA with 25 µL Tris-Cl, pH 8. 2 x 150 bp paired-end sequencing was performed using the Illumina HiSeq4000 system (Nationwide Children’s Hospital Institute for Genomic Medicine). Two independent replicates were performed for each sample, with one replicate consisting of cells prepped from viral infection to sequencing.

### Bioinformatic analyses

For all samples the quality of raw fastq samples was evaluated using FastQC.^56^ Trim Galore!^57^ was then used to trim both ChIP-seq, CUT&RUN, and CUT&Tag reads for adapter sequences and quality. Trimmed reads were aligned to the human genome build hg19/GRCh37 using bowtie2. ChIP-seq reads were aligned with the following parameters (default end-to-end alignment): bowtie2 --no-unal --no-mixed --no-discordant --no-dovetail --phred 33 --q --I 10 --X 1000 --threads 16. CUT&RUN reads were aligned with the following parameters: bowtie2 --no-unal --no-mixed --no-discordant --dovetail --phred 33 --q --I 10 --X 1000 --threads 16. Output SAM files were converted to BAM files, sorted, and indexed using samtools.^58^ Pybedtools^59^ was used to convert BAM files to BED files. To generate bigwig files for visualization, we first converted BED files to spike-in normalized (CUT&RUN and CUT&Tag) or read-count normalized (ChIP-seq) Bedgraph files. We then used the UCSC utility bedGraphToBigWig to generate BigWig files. Replicate samples were verified for high levels (> 0.9 for transcription factors and >0.85 for histone marks) of inter-sample correlation using the UCSC utility wigCorrelate. For CUT&RUN and ChIP-seq, EWS/FLI, LSD1, H3K4me2, and H3K4me3, peaks were called using the default settings of MACS2 callpeak.^60^ For H3K4me1, peaks were called using the --broad setting of callpeak. H3K27ac peaks were called using csaw^61^ using a window of 150 bp, spacing of 50 bp, background signal binned into 2000 bp windows, and a 3-fold increase threshold over global background and an FDR of < 0.05. For CUT&Tag samples (LSD1, H3K27ac and Rb) peaks were called for each sample using the default settings of MACS2 without a control file specified. Then MACS2 bdgdiff with --d1 and --d2 flags used to specify spike-in factors was used to find regions of each sample (LSD1, H3K27ac) with greater signal than Rb, as well as to compare samples in A673, EFKD, and wtEF cells directly. Tracks were generated in the Integrated Genome Browser. ChIPPeakAnno^62^ was used to analyze genomic distribution of peaks and, in concert with bedtools^63^, genome-wide overlaps between groups. HOMER^64^ was used to determine enriched motifs associated with different peaks utilizing the findMotifsGenome.pl script. GSEA^37,65^ was used to analyze functional association between peak-associated genes and EWS/FLI or LSD1 function. deepTools^66^ computeMatrix, plotProfile, and plotHeatmap were used to generate profile and heatmap figures for different groups of binding profiles. Ranked order of super-enhancers (ROSE)^39,40^ was used to identify super-enhancers and super-clusters.

### Quantification and Statistical Analysis

Significance of experimental results was carried out using unpaired t-test for comparing two groups or one-way ANOVA (with multiple comparisons) for comparing three or more groups as appropriate. Significance was determined as a p < 0.05. These statistical tests were performed using GraphPad Prism 8. For GSEA significance was determined using a normalized enrichment score (NES). A result was significant if |NES| > 1.5. HOMER, MACS2 and csaw statistical defaults were used and are described elsewhere.^60,61,64^ For Venn diagram overlaps, p-values were determined using ChIPPeakAnno findOverlapOfPeaks.

### Data and Pipeline Availability

Raw data, bigwigs, and peak calling results are available under the GEO accession: GSE144688

The quality, trimming, and alignment pipelines for single-end ChIP, CUT&RUN, CUT&Tag are available in Singularity containers and can be downloaded from Singularity Hub.

single-end ChIP: shub://ertheisen/southkaibab_centos:hg19v1.centos

CUT&RUN and CUT&Tag: shub://ertheisen/hohriver_centos:hg19v2.centos

Alternatively, the recipe files are available at https://github.com/ertheisen?tab=repositories.

## ACKNOWLEDGMENT

We thank Dr. Ranajeet Saund for technical assistance. This research was supported in part by the High Performance Computing Facility at the Abigail Wexner Research Institute at Nationwide Children’s Hospital.

## SUPPLEMENTAL FIGURES

**Supplementary Figure 1.**
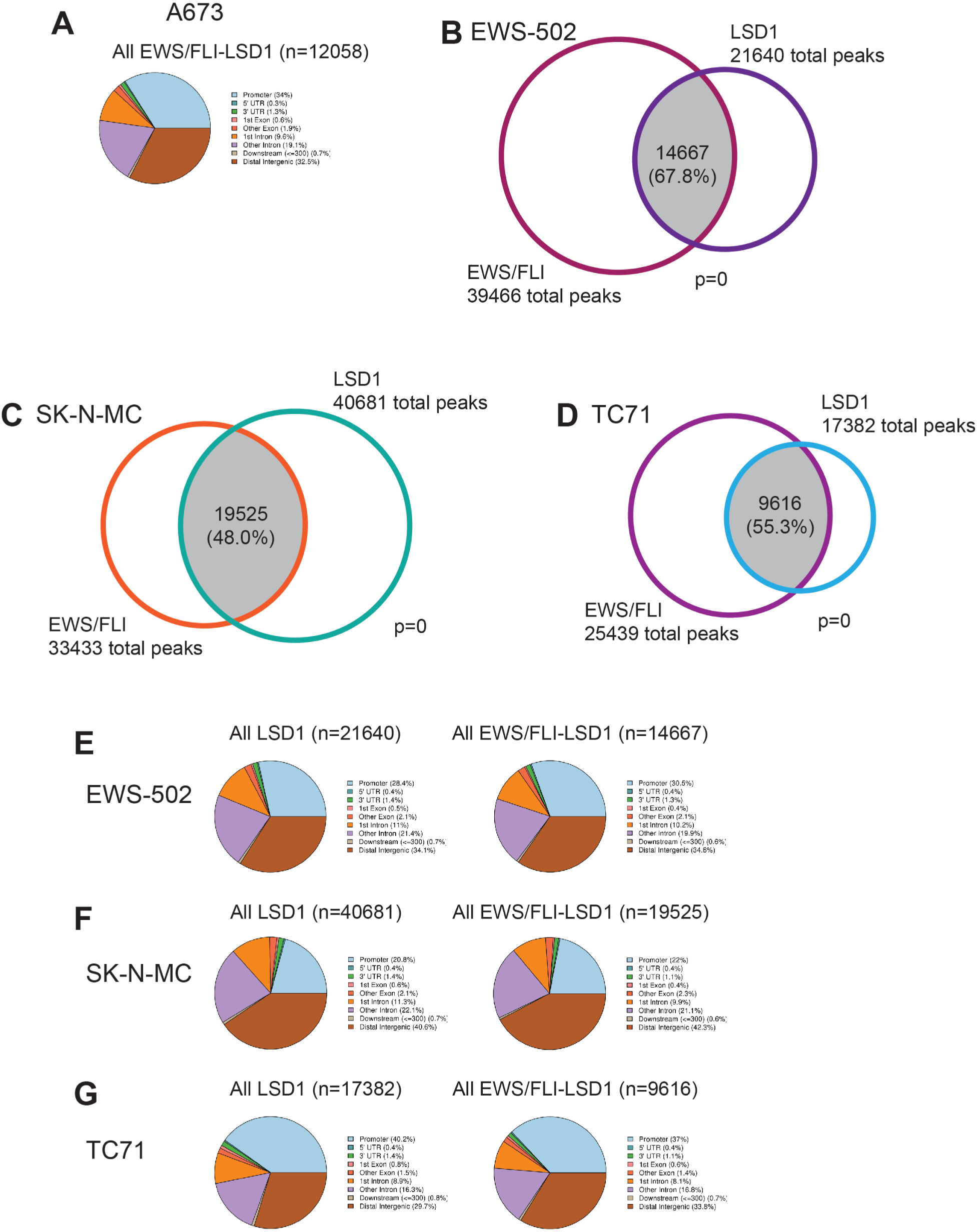
A) Genomic distributions of EWS/FLI-LSD1 coincident peaks in A673 cells. B-D) Venn diagram of EWS/FLI and LSD1 peaks in (B) EWS-502, (C) SK-N-MC, and (D) TC71 cells as determined by ChIPPeakAnno; p-value calculated by ChIPPeakAnno. E-G) Genomic distributions of LSD1 peaks and EWS/FLI-LSD1 coincident peaks in (E) EWS-502, (F) SK-N-MC, and (G) TC71 cells.

**Supplementary Figure 2.**
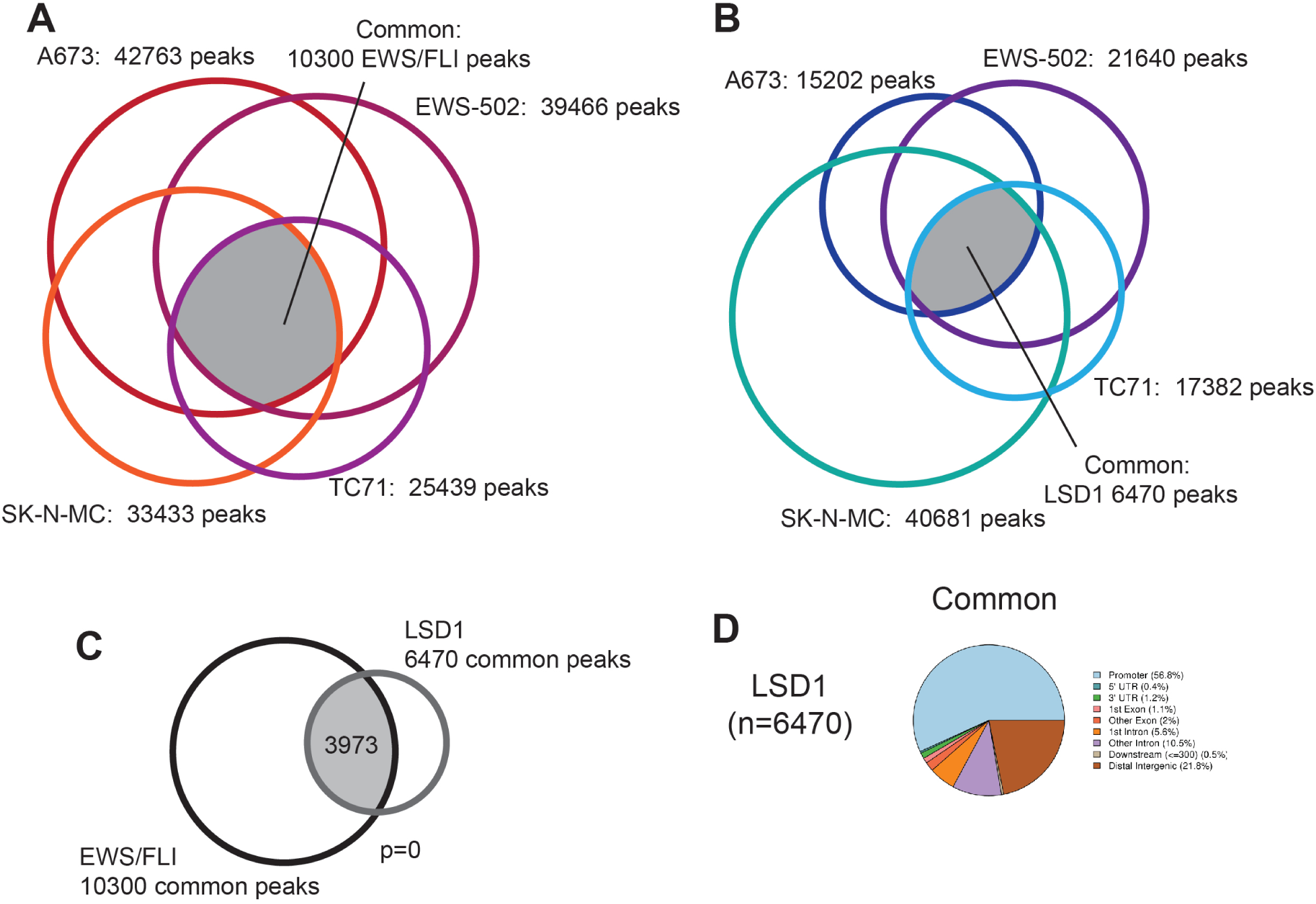
A,B) Venn diagram of (A) EWS/FLI and (B) LSD1 peaks in all cell lines tested as determined by ChIPPeakAnno. C) Venn diagram of EWS/FLI and LSD1 peaks common across all cell lines as determined by ChIPPeakAnno. D) Genomic distributions of LSD1 peaks common across cell lines.

**Supplementary Figure 3.**
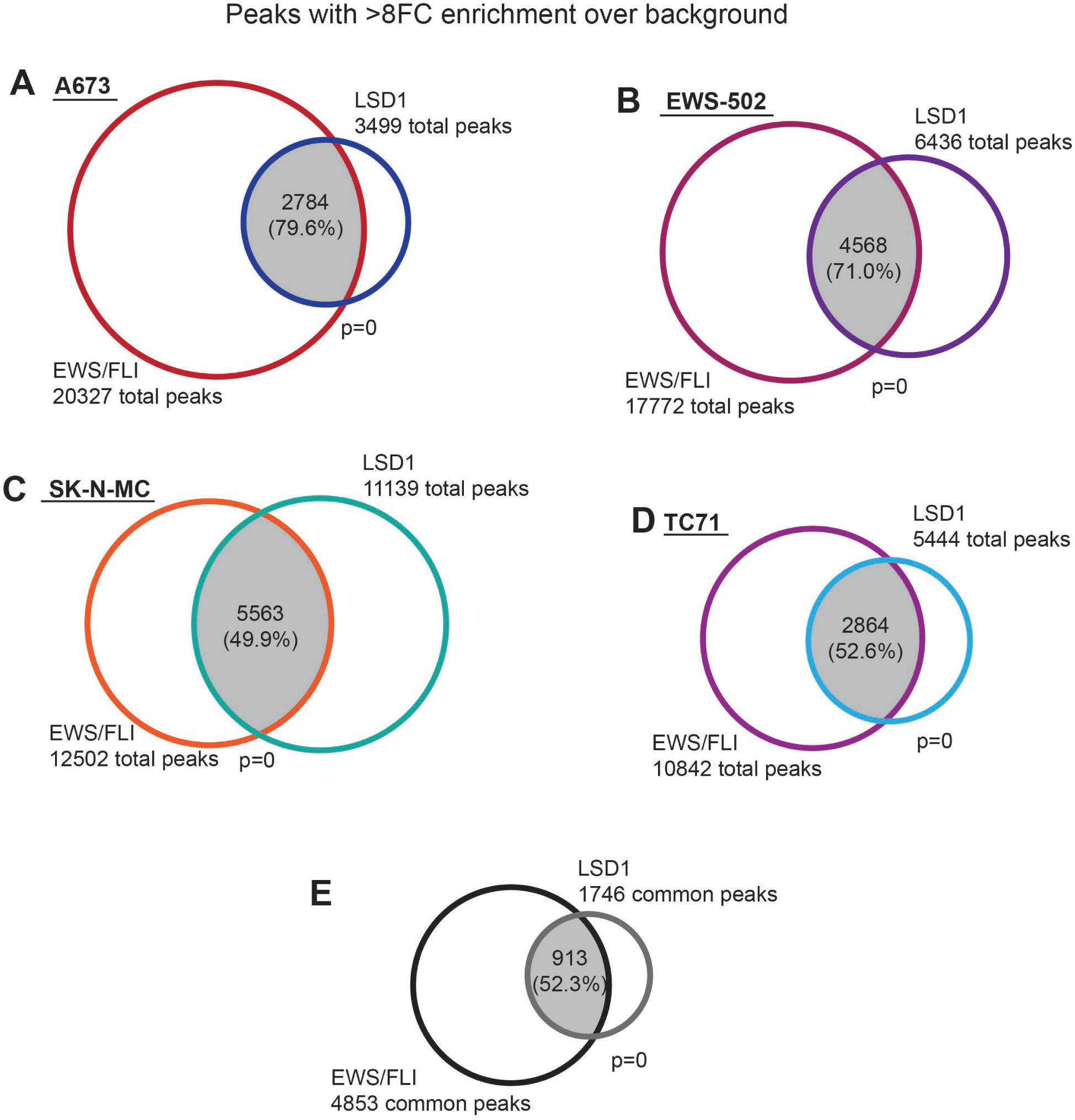
A-E) Venn diagram of EWS/FLI and LSD1 peaks that pass an 8-fold enrichment cutoff in (A) A673 cells, (B) EWS-502 cells, (C) SK-N-MC cells, and (D) TC71 cells as determined by ChIPPeakAnno; p-value calculated by ChIPPeakAnno. (E) shows the overlap between common EWS/FLI and common LSD1 peaks.

**Supplementary Figure 4.**
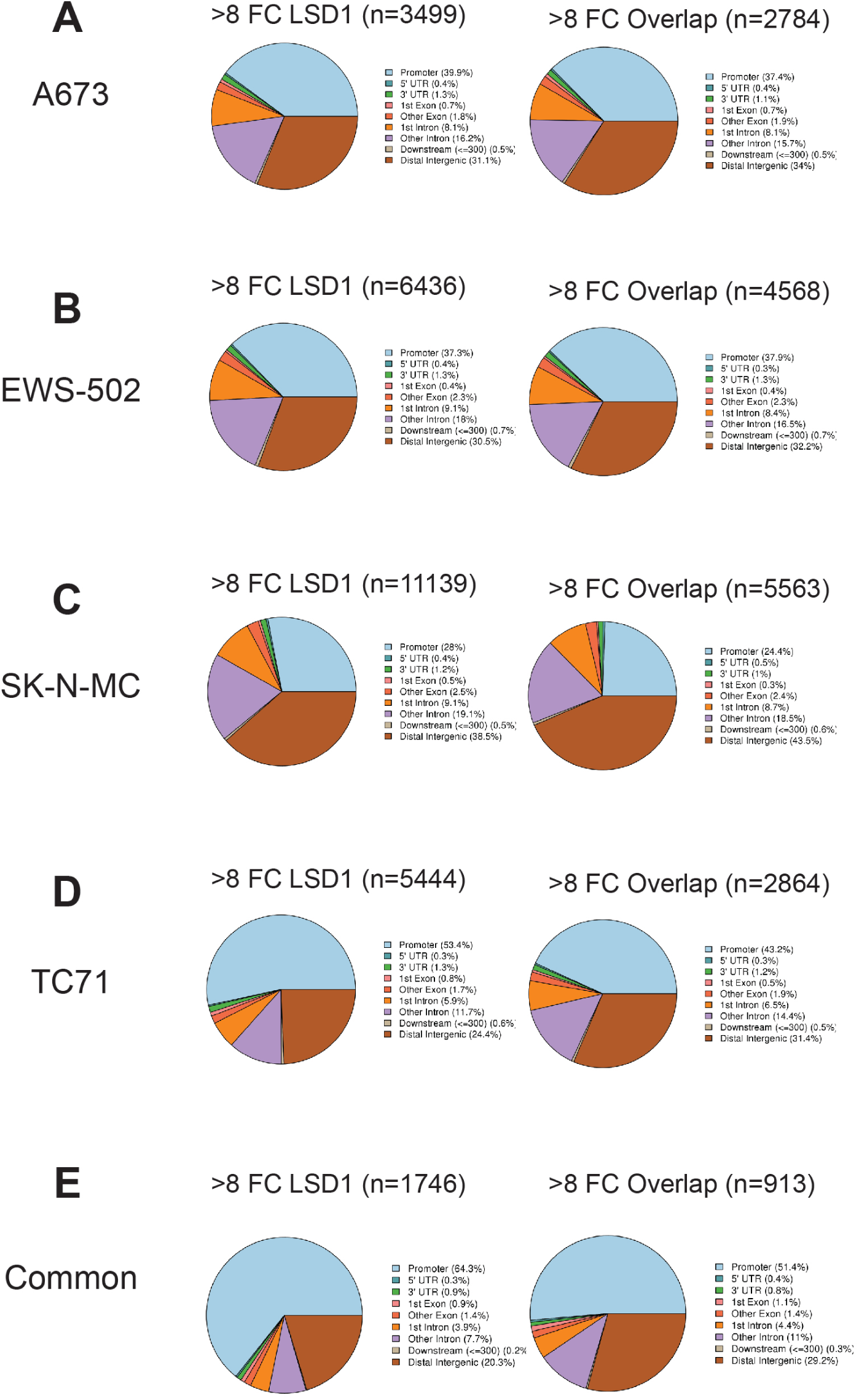
A-E) Genomic distributions of LSD1 peaks and EWS/FLI-LSD1 coincident peaks which pass an 8-fold enrichment in (A) A673, (B) EWS-502, (C) SK-N-MC, and (D) TC71 cells. (E) depicts genomic distribution of peaks common across cell lines.

**Supplementary Figure 5.**
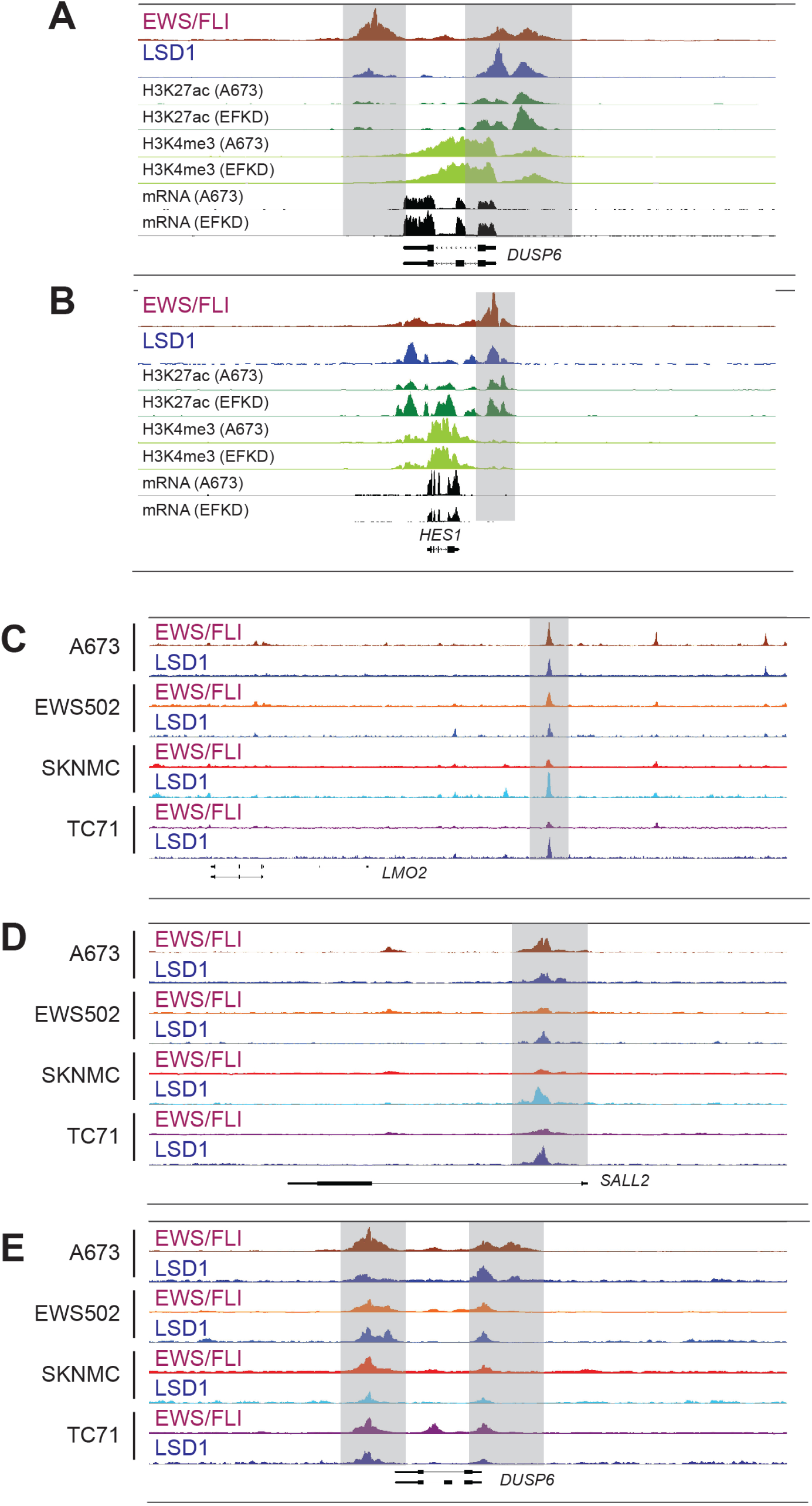
A,B) IGB tracks showing coincidence of EWS/FLI and LSD1 near EWS/FLI-repressed gene *DUSP6* (A) and EWS/FLI-activated gene *HES1* (B). Tracks also show H3K27ac, H3K4me3, and mRNA in the A673 and EWS/FLI-depleted (EFKD) conditions. C-E) Genomic tracks showing localization of LSD1 and EWS/FLI near (C) *LMO2*, (D) *SALL2*, and (E) *DUSP6* in all tested cell lines.

**Supplementary Figure 6.**
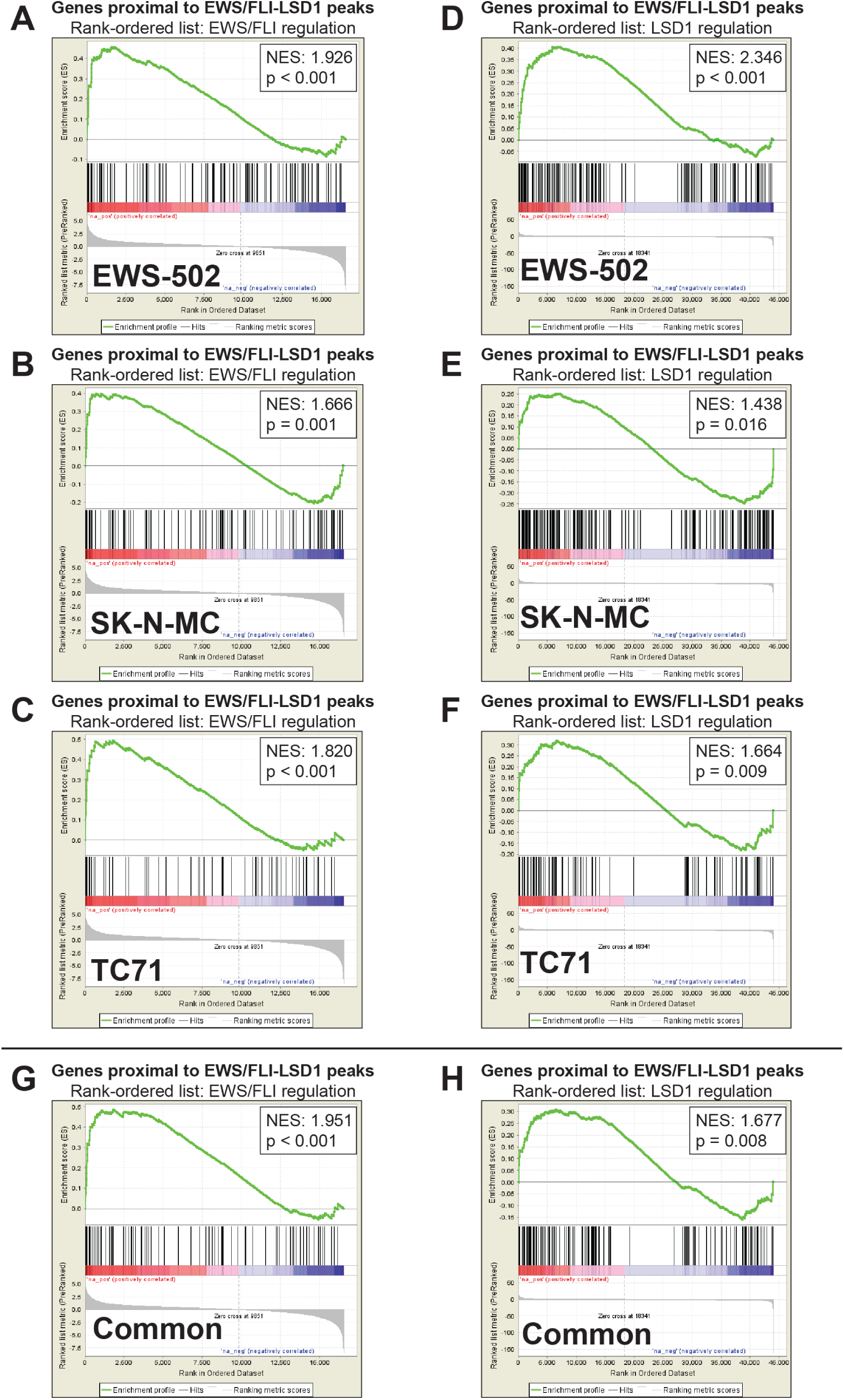
A-H) GSEA results using promoter-proximal EWS/FLI-LSD1 coincident peaks (<5kb to TSS) in different cell lines (A-F) or common to all cell lines (G,H) as the test and (A-C,G) EWS/FLI gene regulation or (D-F,H) LSD1 gene regulation as the rank-ordered dataset. NES=normalized enrichment score. |NES|>1.5 is significant. EWS-502: n=164. SK-N-MC: n=182. TC71: n=87. Common: n=123

**Supplementary Figure 7.**
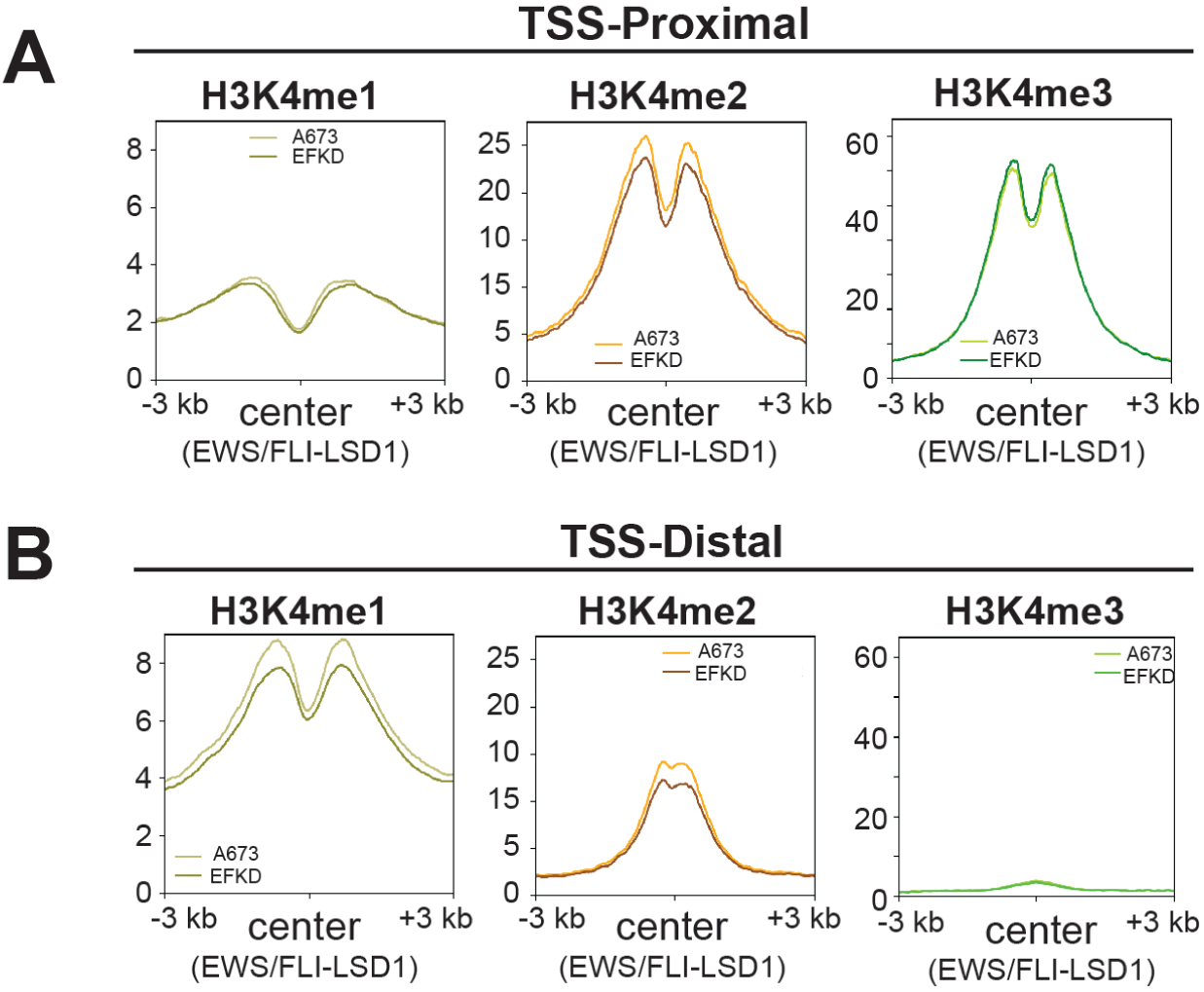
A,B) Profile plots for signal intensity of H3K4me1, H3K4me2, and H3K4me3 within 3 kb of EWS/FLI-LSD1 coincident peaks in either A673 cells or EFKD as specified. Profile plots are separated into those regions either proximal to (A, n=1104) or distal to (B, n=1680) TSS.

**Supplementary Figure 8.**
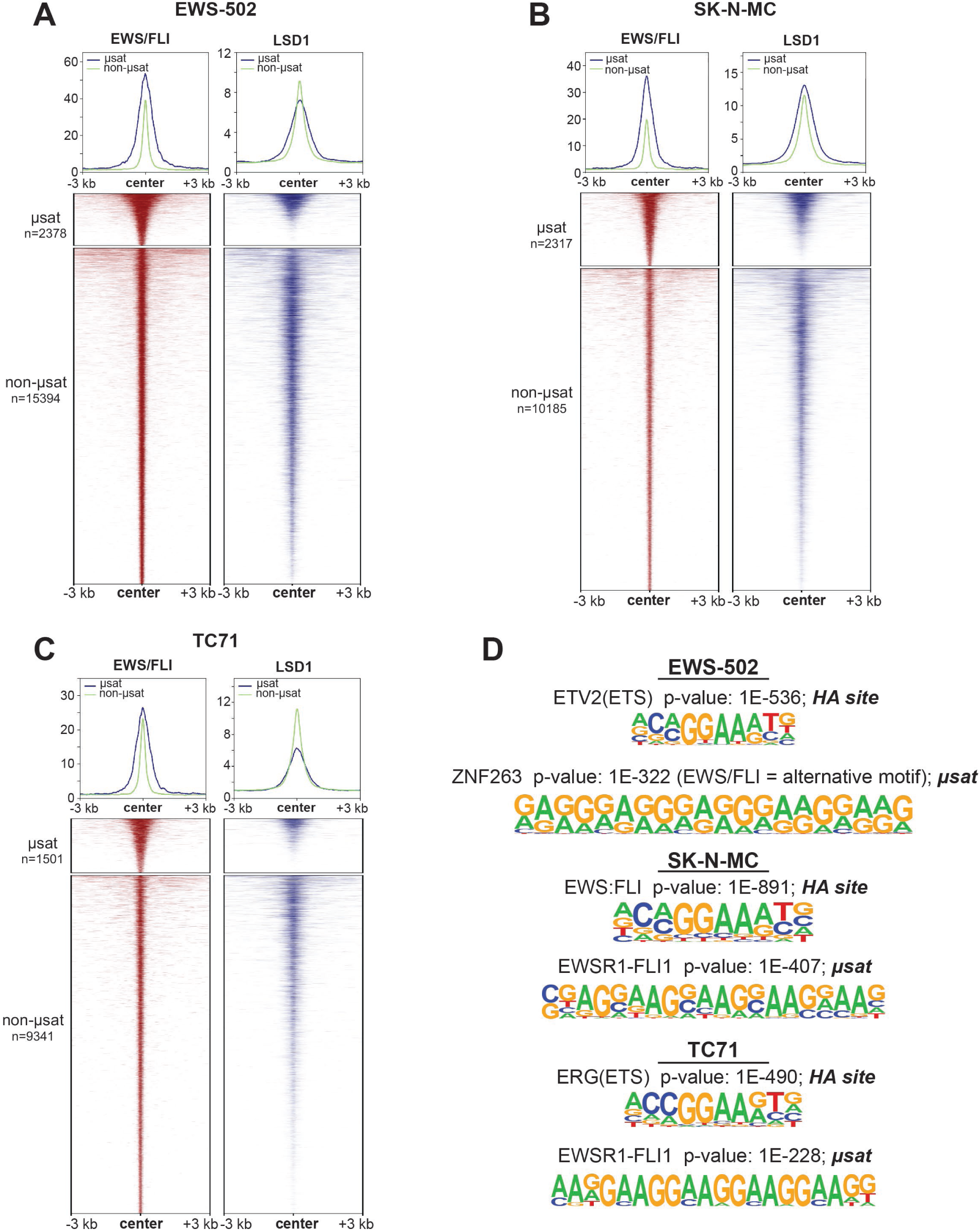
A-C) Profile plots and heatmaps of EWS/FLI (red) and LSD1 (blue) within 3 kb of EWS/FLI peaks in (A) EWS-502, (B) SK-N-MC, and (C) TC71 cells. GGAA-microsatellite (µsat) peaks are represented in profile with a blue line and are the top panel in the heatmap. Non-microsatellite (non-µsat) peaks are represented in profile with a green line and are the bottom panel in the heatmap. D) Top two ranked results from HOMER *de novo* motif enrichment analysis in different cells with significance value.

**Supplementary Figure 9.**
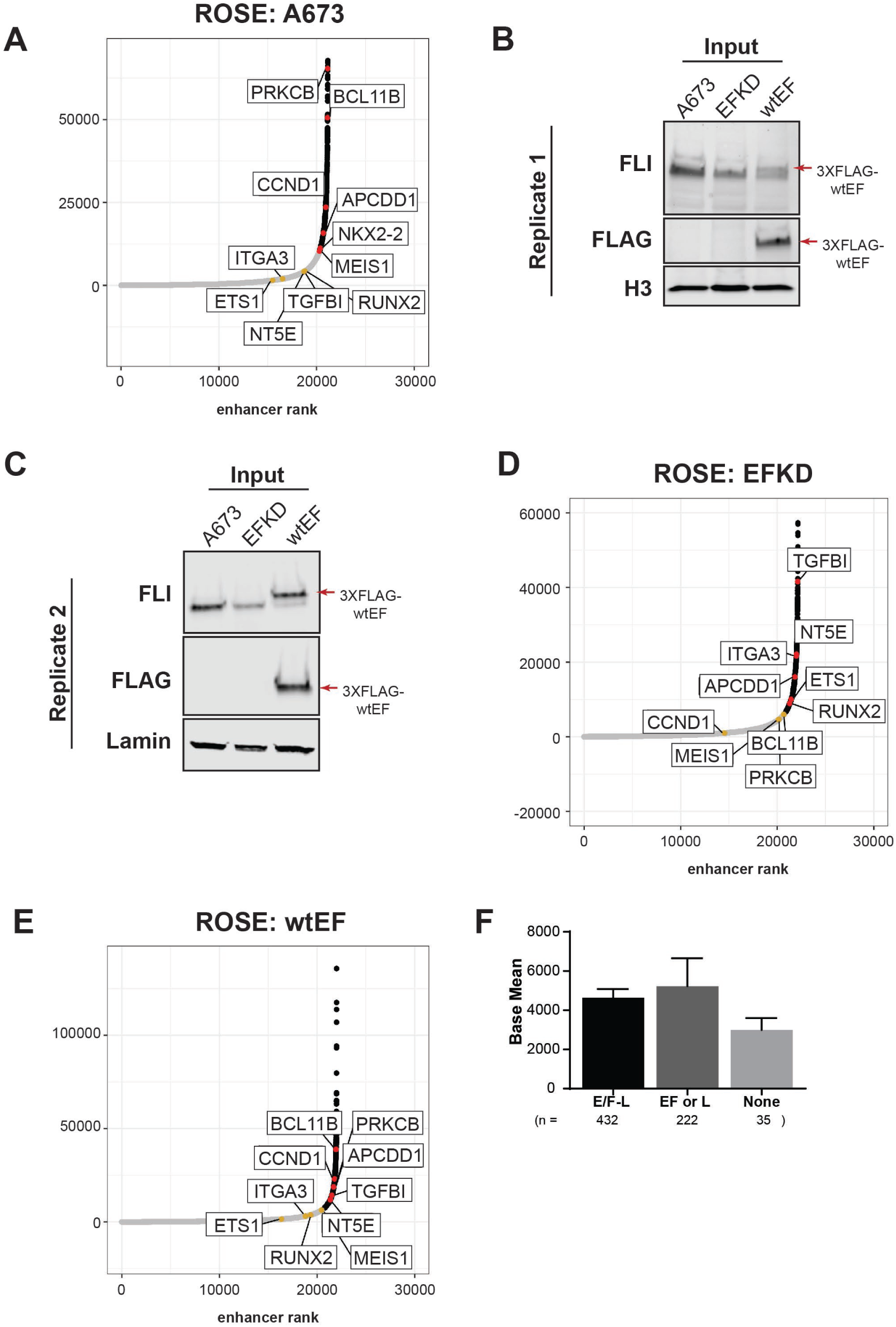
related to Supplementary Tables 1-6. A) Plotted output of the ROSE analysis for super-enhancers in A673 cells. B,C) Western blot validation of EWS/FLI KD in EFKD cells and rescue with 3XFLAG-EWS/FLI in wtEF cells in (B) replicate 1 and (C) replicate 2. Ectopic EWS/FLI was introduced following RNAi-mediated depletion of EWS/FLI using iEF-2 or an iLuc (negative control) construct. D,E) Plotted output of the ROSE analysis for super-enhancers in EFKD and (E) wtEF cells. F) Base mean expression of genes associated with super-enhancers in A673 cells separated by the type of overlap with EWS/FLI and LSD1. E/F-L = EWS/FLI and LSD1, EF = EWS/FLI only, L = LSD1 only, or None. Mean and SD are shown and P-values were determined using one-way ANOVA with multiple comparison testing. No significant differences were detected.

**Supplementary Figure 10.**
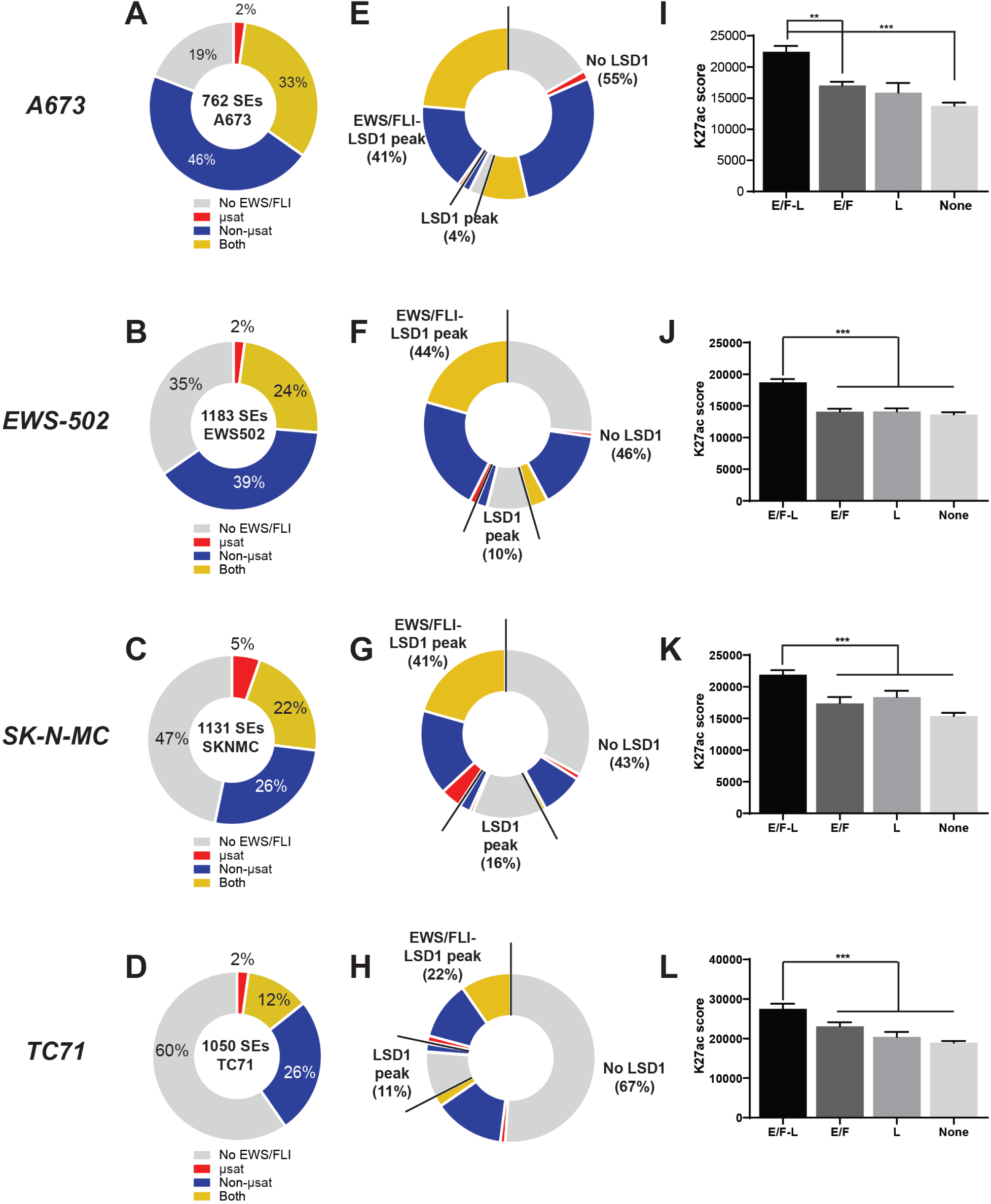
related to Supplementary Tables 7-14. A-D) Pie chart distribution of super-enhancers (SEs) in (A)A673, (B) EWS-502, (C) SK-N-MC, and (D) TC71 cells (CUT&RUN) by type of overlapped EWS/FLI-bound motif. E-H) Pie chart distribution of SEs in (E) A673, (F) EWS-502, (G) SK-N-MC, and (H) TC71 cells (CUT&RUN) by type of EWS/FLI and LSD1 overlap. I-L) H3K27ac score calculated from the ROSE algorithm for SEs in (I) A673, (J) EWS-502, (K) SK-N-MC, and (L) TC71 cells (CUT&RUN) plotted by type of overlap with EWS/FLI and LSD1. E/F-L=EWS/FLI and LSD1 coincident peak, EF=EWS/FLI only, L=LSD1 only. Mean and SD are shown and p-values were determined using one-way ANOVA with multiple comparison testing (***p<0.001, **p<0.01, *p<0.05.) N for differential expression is lower for those K27ac scores because not all genes near SEs were detected by RNA-seq.

**Supplementary Figure 11.**
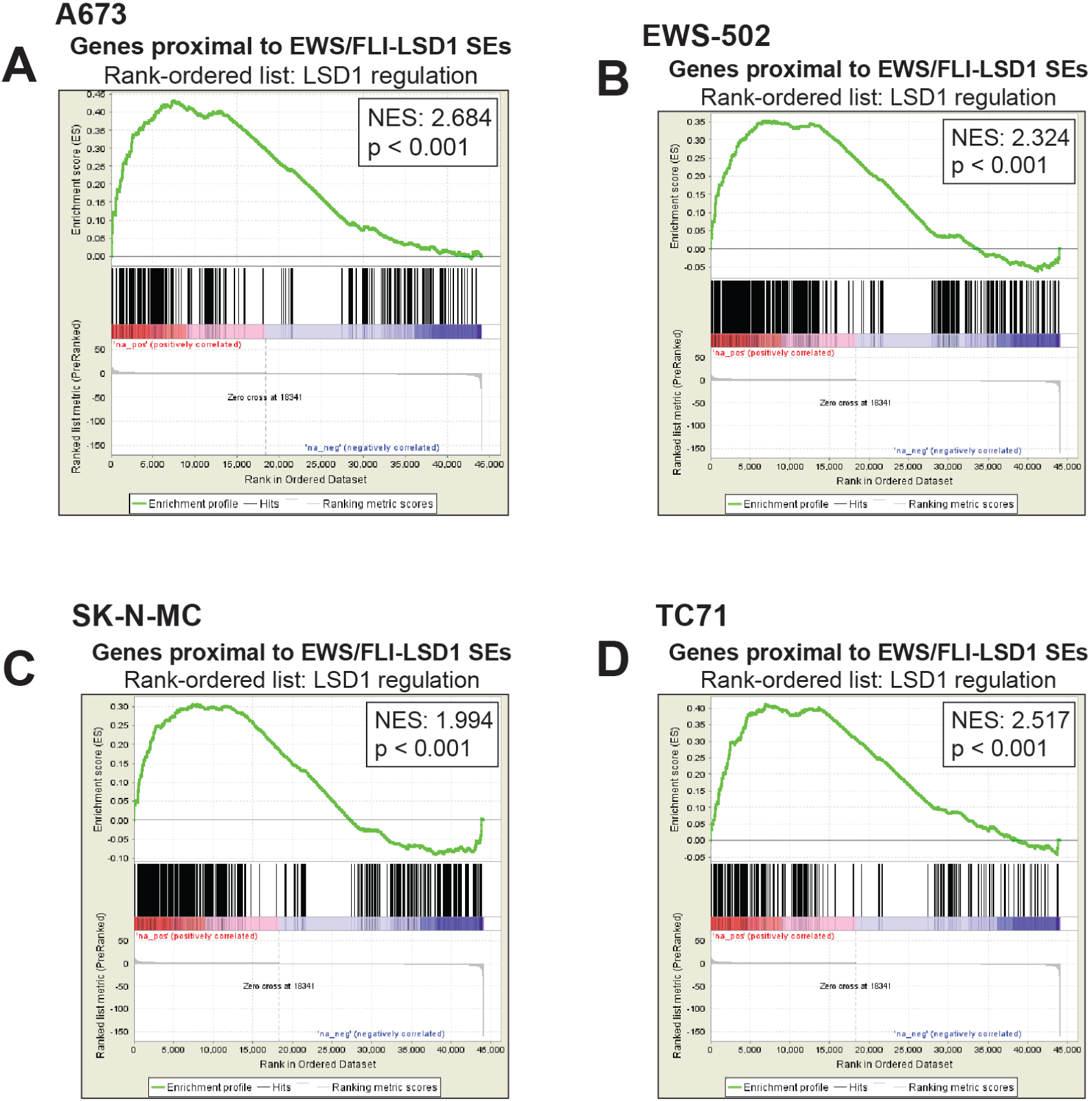
related to Supplementary Tables 8, 10, 12, 14. A-D) GSEA results using genes near SEs with an EWS/FLI-LSD1 coincident peak in (A) A673 cells (CUT&RUN, n=319), (B) EWS-502 cells (n=545), (C) SK-N-MC cells (n=491), and (D) TC71 cells (n=251) as the test set and LSD1 gene regulation in A673 cells as the rank-ordered dataset. NES = normalized enrichment score. |NES| > 1.5 is significant.

**Supplementary Figure 12.**
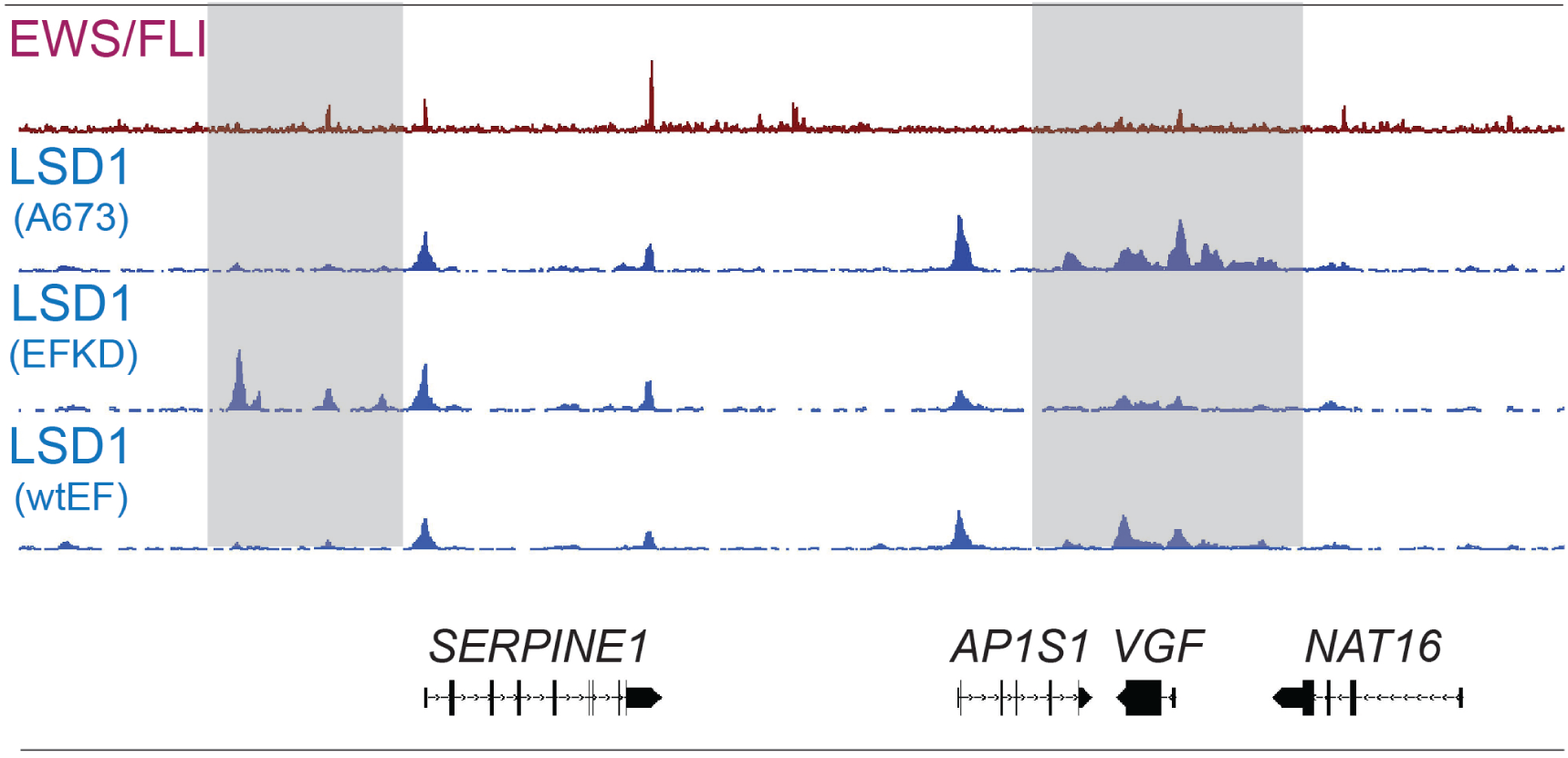
IGB tracks showing EWS/FLI and LSD1 near *SERPINE1, AP1S1, VGF*, and *NAT16*. Tracks show the sensitivity of CUT&Tag to detect changes in LSD1 levels between A673, EFKD, and wtEF cells, with a region over *SERPINE1* showing increased LSD1 with EWS/FLI depletion, and another region near *VGF* showing decreased LSD1 with EWS/FLI depletion.

**Supplementary Figure 13.**
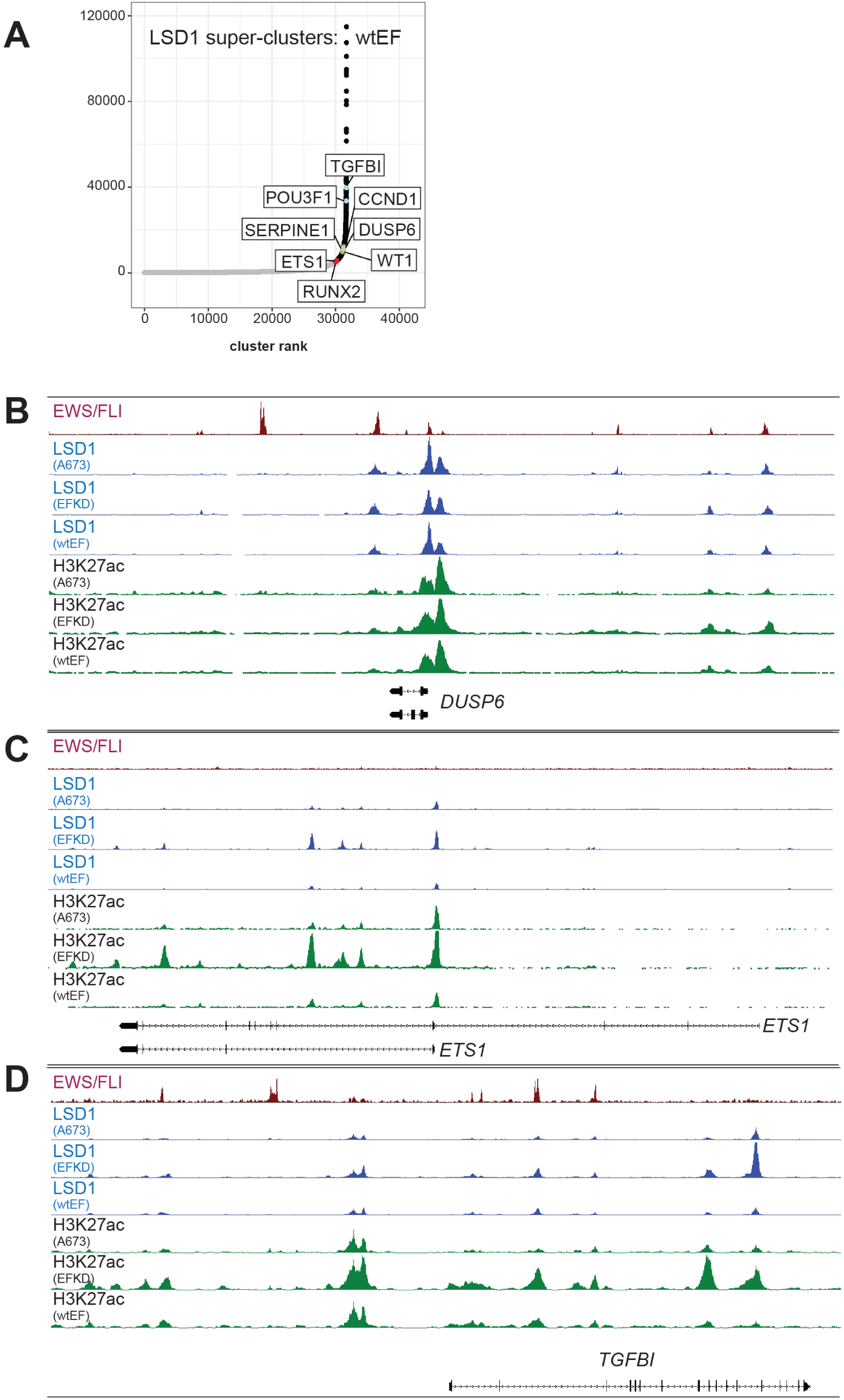
related to Supplementary Tables 1-6, 15-20. A) Plotted output of the ROSE analysis for LSD1 superclusters (SCs) in wtEF cells. B-D) IGB tracks showing coincidence of EWS/FLI, LSD1, and H3K27ac in a super-enhancer and LSD1 super-cluster near *DUSP6* (B), *ETS1* (C), and *TGFBI* (D). Tracks show LSD1 and H3K27ac in A673, EFKD, and wtEF conditions. In (C) the super-enhancer and super-cluster are only present in EFKD cells.

**Supplementary Figure 14.**
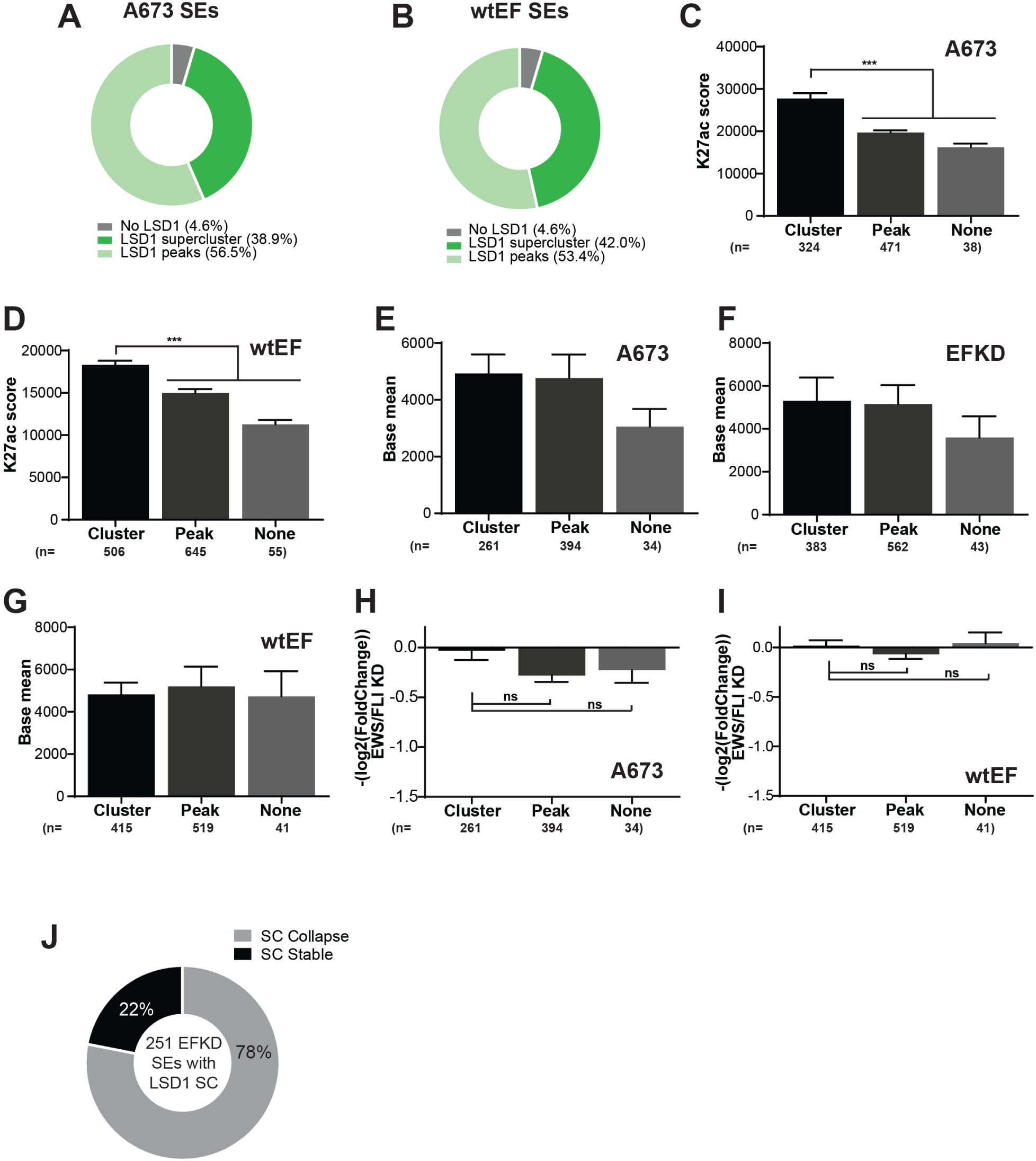
related to Supplementary Tables 2, 4, and 6. A,B) Pie chart distribution showing the overlap of SEs in (A) A673 and (B) wtEF cells with different types of LSD1-binding. C,D) H3K27ac score calculated from the ROSE algorithm for (C) A673 and (D) wtEF cells plotted by type of overlap with LSD1. E-G) Base mean expression of genes associated with super-enhancers in (E) A673, (F) EFKD, and (G) wtEF cells separated by the type of overlap with LSD1. H,I) EWS/FLI-mediated differential expression of genes near SEs in (H) A673 and (I) wtEF cells plotted by type of overlap with LSD1. Mean and SD are shown and p-values were determined using one-way ANOVA with multiple comparison testing (***p<0.001, **p<0.01, *p< 0.05). N for differential expression and base mean is lower for those K27ac scores because not all genes near SEs were detected by RNA-seq. J) Pie chart distribution showing the number of LSD1 SCs in LSD1 SC-containing SEs unique to EFKD cells that collapse with EWS/FLI expression.

## SUPPLEMENTARY FILES

### Supplementary_File_1.xlsx

Contains Supplementary Tables 1-20.

Supplementary Tables 1-2: 1) Results of the ROSE analysis for superenhancers in A673 cells and 2) annotated A673 superenhancers. The K27ac data used was generated by CUT&Tag.

Supplementary Tables 3-4: 3) Results of the ROSE analysis for superenhancers in EFKD cells and 4) annotated EFKD superenhancers. The K27ac data used was generated by CUT&Tag.

Supplementary Tables 5-6: 5) Results of the ROSE analysis for superenhancers in wtEF cells and 6) annotated wtEF superenhancers. The K27ac data used was generated by CUT&Tag.

Supplementary Tables 7-8: 7) Results of the ROSE analysis for superenhancers in A673 cells and 8) annotated A673 superenhancers. The K27ac data used was generated by CUT&RUN.

Supplementary Tables 9-10: 9) Results of the ROSE analysis for superenhancers in EWS-502 cells and 10) annotated EWS-502 superenhancers. The K27ac data used was generated by CUT&RUN.

Supplementary Tables 11-12: 11) Results of the ROSE analysis for superenhancers in SK-N-MC cells and 12) annotated SK-N-MC superenhancers. The K27ac data used was generated by CUT&RUN.

Supplementary Tables 13-14: 13) Results of the ROSE analysis for superenhancers in TC71 cells and 14) annotated TC71 superenhancers. The K27ac data used was generated by CUT&RUN.

Supplementary Tables 15-16: 15) Results of the ROSE analysis for LSD1 superclusters in A673 cells and 16) annotated A673 LSD1 superclusters. The LSD1 data used was generated by CUT&Tag.

Supplementary Tables 17-18: 17) Results of the ROSE analysis for LSD1 superclusters in EFKD cells and 18) annotated EFKD LSD1 superclusters. The LSD1 data used was generated by CUT&Tag.

Supplementary Tables 19-20: 19) Results of the ROSE analysis for LSD1 superclusters in wtEF cells and 20) annotated wtEF LSD1 superclusters. The LSD1 data used was generated by CUT&Tag.

### Supplementary_Table_21.xlsx

Contains a list of key resources for the work performed in this report.

## REFERENCES

1. Burningham, Z., Hashibe, M., Spector, L. & Schiffman, J. D. The Epidemiology of Sarcoma. Clin. Sarcoma Res. 2, 14 (2012).

2. Delattre, O. et al. Gene fusion with an ETS DNA-binding domain caused by chromosome translocation in human tumours. Nature 359, 162–165 (1992).

3. Turc-Carel, C., Philip, I., Berger, M. P., Philip, T. & Lenoir, G. M. Chromosome study of Ewing’s sarcoma (ES) cell lines. Consistency of a reciprocal translocation t(11;22)(q24;q12). Cancer Genet. Cytogenet. 12, 1–19 (1984).

4. Turc-Carel, C. et al. Chromosomes in Ewing’s sarcoma. I. An evaluation of 85 cases of remarkable consistency of t(11;22)(q24;q12). Cancer Genet. Cytogenet. 32, 229–238 (1988).

5. Boulay, G. et al. Cancer-Specific Retargeting of BAF Complexes by a Prion-like Domain. Cell 171, 163-178.e19 (2017).

6. May, W. A. et al. Ewing sarcoma 11;22 translocation produces a chimeric transcription factor that requires the DNA-binding domain encoded by FLI1 for transformation. Proc. Natl. Acad. Sci. U. S. A. 90, 5752–5756 (1993).

7. May, W. A. et al. The Ewing’s sarcoma EWS/FLI-1 fusion gene encodes a more potent transcriptional activator and is a more powerful transforming gene than FLI-1. Mol. Cell. Biol. 13, 7393–7398 (1993).

8. Riggi, N. et al. EWS-FLI1 Utilizes Divergent Chromatin Remodeling Mechanisms to Directly Activate or Repress Enhancer Elements in Ewing Sarcoma. Cancer Cell 26, 668–681 (2014).

9. Sankar, S. et al. Mechanism and relevance of EWS/FLI-mediated transcriptional repression in Ewing sarcoma. Oncogene 32, 5089–5100 (2013).

10. Smith, R. et al. Expression profiling of EWS/FLI identifies NKX2.2 as a critical target gene in Ewing’s sarcoma. Cancer Cell 9, 405–416 (2006).

11. Tomazou, E. M. et al. Epigenome Mapping Reveals Distinct Modes of Gene Regulation and Widespread Enhancer Reprogramming by the Oncogenic Fusion Protein EWS-FLI1. Cell Rep. 10, 1082–1095 (2015).

12. Gangwal, K., Close, D., Enriquez, C. A., Hill, C. P. & Lessnick, S. L. Emergent Properties of EWS/FLI Regulation via GGAA Microsatellites in Ewing’s Sarcoma. Genes Cancer 1, 177–187 (2010).

13. Gangwal, K. et al. Microsatellites as EWS/FLI response elements in Ewing’s sarcoma. Proc. Natl. Acad. Sci. 105, 10149–10154 (2008).

14. Guillon, N. et al. The Oncogenic EWS-FLI1 Protein Binds In Vivo GGAA Microsatellite Sequences with Potential Transcriptional Activation Function. PLoS ONE 4, e4932 (2009).

15. Kwon, I. et al. Phosphorylation-Regulated Binding of RNA Polymerase II to Fibrous Polymers of Low-Complexity Domains. Cell 155, 1049–1060 (2013).

16. Theisen, E. R. et al. Transcriptomic analysis functionally maps the intrinsically disordered domain of EWS/FLI and reveals novel transcriptional dependencies for oncogenesis. Genes Cancer 10, 21–38 (2019).

17. Sankar, S. et al. Reversible LSD1 Inhibition Interferes with Global EWS/ETS Transcriptional Activity and Impedes Ewing Sarcoma Tumor Growth. Clin. Cancer Res. 20, 4584–4597 (2014).

18. Shi, Y. et al. Histone Demethylation Mediated by the Nuclear Amine Oxidase Homolog LSD1. Cell 119, 941–953 (2004).

19. Metzger, E. et al. LSD1 demethylates repressive histone marks to promote androgen-receptor-dependent transcription. Nature 437, 436–439 (2005).

20. Huang, J. et al. p53 is regulated by the lysine demethylase LSD1. Nature 449, 105–108 (2007).

21. Wang, J. et al. The lysine demethylase LSD1 (KDM1) is required for maintenance of global DNA methylation. Nat. Genet. 41, 125–129 (2009).

22. Yang, M. et al. Structural Basis for CoREST-Dependent Demethylation of Nucleosomes by the Human LSD1 Histone Demethylase. Mol. Cell 23, 377–387 (2006).

23. Whyte, W. A. et al. Enhancer decommissioning by LSD1 during embryonic stem cell differentiation. Nature 482, 221–225 (2012).

24. Hatzi, K. et al. Histone demethylase LSD1 is required for germinal center formation and BCL6-driven lymphomagenesis. Nat. Immunol. 20, 86 (2019).

25. Maiques-Diaz, A. et al. Enhancer Activation by Pharmacologic Displacement of LSD1 from GFI1 Induces Differentiation in Acute Myeloid Leukemia. Cell Rep. 22, 3641–3659 (2018).

26. Sugino, N. et al. A novel LSD1 inhibitor NCD38 ameliorates MDS-related leukemia with complex karyotype by attenuating leukemia programs via activating super-enhancers. Leukemia 31, 2303–2314 (2017).

27. Harris, W. J. et al. The histone demethylase KDM1A sustains the oncogenic potential of MLL-AF9 leukemia stem cells. Cancer Cell 21, 473–487 (2012).

28. Hayami, S. et al. Overexpression of LSD1 contributes to human carcinogenesis through chromatin regulation in various cancers. Int. J. Cancer 128, 574–586 (2011).

29. Kahl, P. et al. Androgen receptor coactivators lysine-specific histone demethylase 1 and four and a half LIM domain protein 2 predict risk of prostate cancer recurrence. Cancer Res. 66, 11341–11347 (2006).

30. Lim, S. et al. Lysine-specific demethylase 1 (LSD1) is highly expressed in ER-negative breast cancers and a biomarker predicting aggressive biology. Carcinogenesis 31, 512–520 (2010).

31. Schulte, J. H. et al. Lysine-specific demethylase 1 is strongly expressed in poorly differentiated neuroblastoma: implications for therapy. Cancer Res. 69, 2065–2071 (2009).

32. Bennani-Baiti, I. M., Machado, I., Llombart-Bosch, A. & Kovar, H. Lysine-specific demethylase 1 (LSD1/KDM1A/AOF2/BHC110) is expressed and is an epigenetic drug target in chondrosarcoma, Ewing’s sarcoma, osteosarcoma, and rhabdomyosarcoma. Hum. Pathol. 43, 1300–1307 (2012).

33. Schildhaus, H.-U. et al. Lysine-specific demethylase 1 is highly expressed in solitary fibrous tumors, synovial sarcomas, rhabdomyosarcomas, desmoplastic small round cell tumors, and malignant peripheral nerve sheath tumors. Hum. Pathol. 42, 1667–1675 (2011).

34. Pishas, K. I. et al. Therapeutic Targeting of KDM1A/LSD1 in Ewing Sarcoma with SP-2509 Engages the Endoplasmic Reticulum Stress Response. Mol. Cancer Ther. 17, 1902–1916 (2018).

35. Zhao, Z.-K. et al. Overexpression of lysine specific demethylase 1 predicts worse prognosis in primary hepatocellular carcinoma patients. World J. Gastroenterol. 18, 6651–6656 (2012).

36. Sankar, S. et al. EWS and RE1-Silencing Transcription Factor Inhibit Neuronal Phenotype Development and Oncogenic Transformation in Ewing Sarcoma. Genes Cancer 4, 213–223 (2013).

37. Subramanian, A. et al. Gene set enrichment analysis: A knowledge-based approach for interpreting genome-wide expression profiles. Proc. Natl. Acad. Sci. 102, 15545–15550 (2005).

38. Boulay, G. et al. Epigenome editing of microsatellite repeats defines tumor-specific enhancer functions and dependencies. Genes Dev. (2018) doi: 10.1101/gad.315192.118.

39. Lovén, J. et al. Selective Inhibition of Tumor Oncogenes by Disruption of Super-Enhancers. Cell 153, 320–334 (2013).

40. Whyte, W. A. et al. Master Transcription Factors and Mediator Establish Super-Enhancers at Key Cell Identity Genes. Cell 153, 307–319 (2013).

41. Kennedy, A. L. et al. Functional, chemical genomic, and super-enhancer screening identify sensitivity to cyclin D1/CDK4 pathway inhibition in Ewing sarcoma. Oncotarget 6, 30178–30193 (2015).

42. Johnson, K. M. et al. Role for the EWS domain of EWS/FLI in binding GGAA-microsatellites required for Ewing sarcoma anchorage independent growth. Proc. Natl. Acad. Sci. U. S. A. 114, 9870–9875 (2017).

43. Hnisz, D. et al. Super-Enhancers in the Control of Cell Identity and Disease. Cell 155, 934–947 (2013).

44. Agarwal, S. et al. LSD1/KDM1A Maintains Genome-wide Homeostasis of Transcriptional Enhancers. bioRxiv 146357 (2017) doi: 10.1101/146357.

45. Kearns, N. A. et al. Functional annotation of native enhancers with a Cas9-histone demethylase fusion. Nat. Methods 12, 401–403 (2015).

46. Mendenhall, E. M. et al. Locus-specific editing of histone modifications at endogenous enhancers. Nat. Biotechnol. 31, 1133–1136 (2013).

47. Lee, C. et al. Lsd1 as a therapeutic target in Gfi1-activated medulloblastoma. Nat. Commun. 10, 332 (2019).

48. Cai, C. et al. Lysine-Specific Demethylase 1 Has Dual Functions as a Major Regulator of Androgen Receptor Transcriptional Activity. Cell Rep. 9, 1618–1627 (2014).

49. Sehrawat, A. et al. LSD1 activates a lethal prostate cancer gene network independently of its demethylase function. Proc. Natl. Acad. Sci. U. S. A. 115, E4179–E4188 (2018).

50. Melot, T. et al. Production and characterization of mouse monoclonal antibodies to wild-type and oncogenic FLI-1 proteins. Hybridoma 16, 457–464 (1997).

51. Lessnick, S. L., Dacwag, C. S. & Golub, T. R. The Ewing’s sarcoma oncoprotein EWS/FLI induces a p53-dependent growth arrest in primary human fibroblasts. Cancer Cell 1, 393–401 (2002).

52. Sankar, S. et al. A Novel Role for Keratin 17 in Coordinating Oncogenic Transformation and Cellular Adhesion in Ewing Sarcoma. Mol. Cell. Biol. 33, 4448–4460 (2013).

53. Kaya-Okur, H. S. et al. CUT&Tag for efficient epigenomic profiling of small samples and single cells. Nat. Commun. 10, 1–10 (2019).

54. Skene, P. J. & Henikoff, S. An efficient targeted nuclease strategy for high-resolution mapping of DNA binding sites. eLife 6, (2017).

55. Buenrostro, J. D. et al. Single-cell chromatin accessibility reveals principles of regulatory variation. Nature 523, 486–490 (2015).

56. Andrews, S. FastQC: a quality control tool for high throughput sequence data.

57. Krueger, F. Trim Galore!: A wrapper tool around Cutadapt and FastQC to consistently apply quality and adapter trimming to FastQ files.

58. Tools (written in C using htslib) for manipulating next-generation sequencing data: samtools/samtools. (samtools, 2019).

59. Dale, R. K., Pedersen, B. S. & Quinlan, A. R. Pybedtools: a flexible Python library for manipulating genomic datasets and annotations. Bioinformatics 27, 3423–3424 (2011).

60. Feng, J., Liu, T., Qin, B., Zhang, Y. & Liu, X. S. Identifying ChIP-seq enrichment using MACS. Nat. Protoc. 7, (2012).

61. Lun, A. T. L. & Smyth, G. K. csaw: a Bioconductor package for differential binding analysis of ChIP-seq data using sliding windows. Nucleic Acids Res. 44, e45 (2016).

62. Zhu, L. J. et al. ChIPpeakAnno: a Bioconductor package to annotate ChIP-seq and ChIP-chip data. BMC Bioinformatics 11, 237 (2010).

63. Quinlan, A. R. & Hall, I. M. BEDTools: a flexible suite of utilities for comparing genomic features. Bioinformatics 26, 841–842 (2010).

64. Heinz, S. et al. Simple Combinations of Lineage-Determining Transcription Factors Prime cis-Regulatory Elements Required for Macrophage and B Cell Identities. Mol. Cell 38, 576–589 (2010).

65. Mootha, V. K. et al. PGC-1α-responsive genes involved in oxidative phosphorylation are coordinately downregulated in human diabetes. Nat. Genet. 34, 267 (2003).

66. Ramírez, F. et al. deepTools2: a next generation web server for deep-sequencing data analysis. Nucleic Acids Res. 44, W160–W165 (2016).

